# Elucidation and engineering of arabinofuranosyltransferase to enable total *de novo* biosynthesis of paris saponins in yeast

**DOI:** 10.64898/2025.12.07.692791

**Authors:** Haowen Wang, Yuxin Yang, Ziya Wu, Huan Zhao, Yinying Ba, Chi Zhang, Chenxing Sun, Zihan Yu, Bowen Qiu, Xuan Liu, Yating Hu, Xianan Zhang

**Author notes:** contributed equally to this work (Haowen Wang, Yuxin Yang, Ziya Wu). Corresponding Authors; (Xianan Zhang). (Yating Hu).

## Abstract

Paris saponins (PSs) are structurally complex steroidal saponins that, due to their diverse glycosylation patterns, exhibit a range of significant pharmacological activities, including anti-tumor and antibacterial effects. However, incomplete characterization of the key enzymes responsible for glycosylation modifications has hindered their efficient heterologous biosynthesis. In this study, we reprogrammed the sugar donor specificity of a steroidal rhamnosyltransferase (UGT93M3) to enable the transfer of arabinofuranose (Ara*f*). Through structural analysis, we identified key amino acid residues (368H/Q) that play an important role in determining Ara*f* donor specificity. Guided by this insight, we successfully reconstructed the paris saponin I (PSI) biosynthetic pathway in *Saccharomyces cerevisiae* using engineered enzymes. To address challenges related to donor availability, we introduced UDP-sugar biosynthetic modules (UDP-Rha and UDP-Ara*f*) into yeast. With this integrated platform, we were able to *de novo* produce a range of paris saponins, including diosgenin-3-*O*-glucosyl-(1→6)-glucoside (DGG), diosgenin-3-*O*-rhamnosyl(1→2) [glucosyl(1→6)]glucoside (DRGG) and paris saponin II. This work establishes a novel microbial platform for the sustainable production of paris saponins, particularly PSI, advancing the biosynthesis of steroidal glycosides and providing a potential strategy for the industrial-scale production of bioactive saponins.

**Teaser:** In this study, we report the first complete *de novo* biosynthesis of four bioactive PSs, PSI, PSII, DGG and DRGG, in *Saccharomyces cerevisiae* from simple carbon source. Key advances include: **(i)** A single amino-acid switch (N368H) in the rhamnosyltransferase UGT93M3 endowed high-efficiency transfer of the rare five-membered arabinofuranose, solving a bottleneck in PS I biosynthesis; **(ii)** Elucidation of the molecular basis for sugar-donor specificity through AlphaFold3 docking and 300-ns molecular-dynamics simulations, revealing a histidine “latch” that stabilizes UDP-Ara*f* in the catalytic pose; **(iii)** Construction of a 16-gene yeast chassis that integrates plant P450s, optimized glycosyltransferases, and *de novo* modules for UDP-rhamnose and UDP-arabinofuranose supply, achieving *de novo* microbial production of PS II, DGG, DRGG and PS I from glucose alone.

## 1. Introduction

Paris saponins are isospirostan-type steroid saponins, predominantly glycosylated at the C3-OH position with sugar moieties consisting of variable numbers of glucose, rhamnose, and arabinose units(Figure 1). They predominantly are found in plant species of the genus *Paris* and the genus *Trillium* within the family Liliaceae, including *Paris polyphylla* var. *yunnanensis* (*Ppy*), *Paris polyphylla* var. *chinensis* (*Ppc*), and *Trillium tschonoskii* Maxim (*Tt*). These compounds exhibit diverse pharmacological activities, including antitumor, antimicrobial and anti-inflammatory effects(*1–3*). Among them, paris saponin I, also known as polyohyllin D, stands out as one of the earliest purified bioactive components from *P. polyphylla*. It exerts cytotoxic effects on various tumor cells(*4*), notably hepatocellular carcinoma and triple-negative breast cancer, by primarily inducing mitochondrial-mediated apoptosis and suppressing cell migration(*5, 6*). More recently, paris saponin I has also been shown to reverse drug resistance in advanced non-small-cell lung cancer to frontline therapeutics such as osimertinib and gefitinib(*7, 8*). Nevertheless, the sustainable supply of paris saponins faces serious limitations due to the endangered status of their botanical sources and protracted growth cycles (typically 7–8 years)(*9*). Additionally, the chemical synthesis of these saponins entails numerous intricate steps and generally proceeds in low yield, posing substantial barriers to their clinical advancement and practical use.

**Figure 1.**
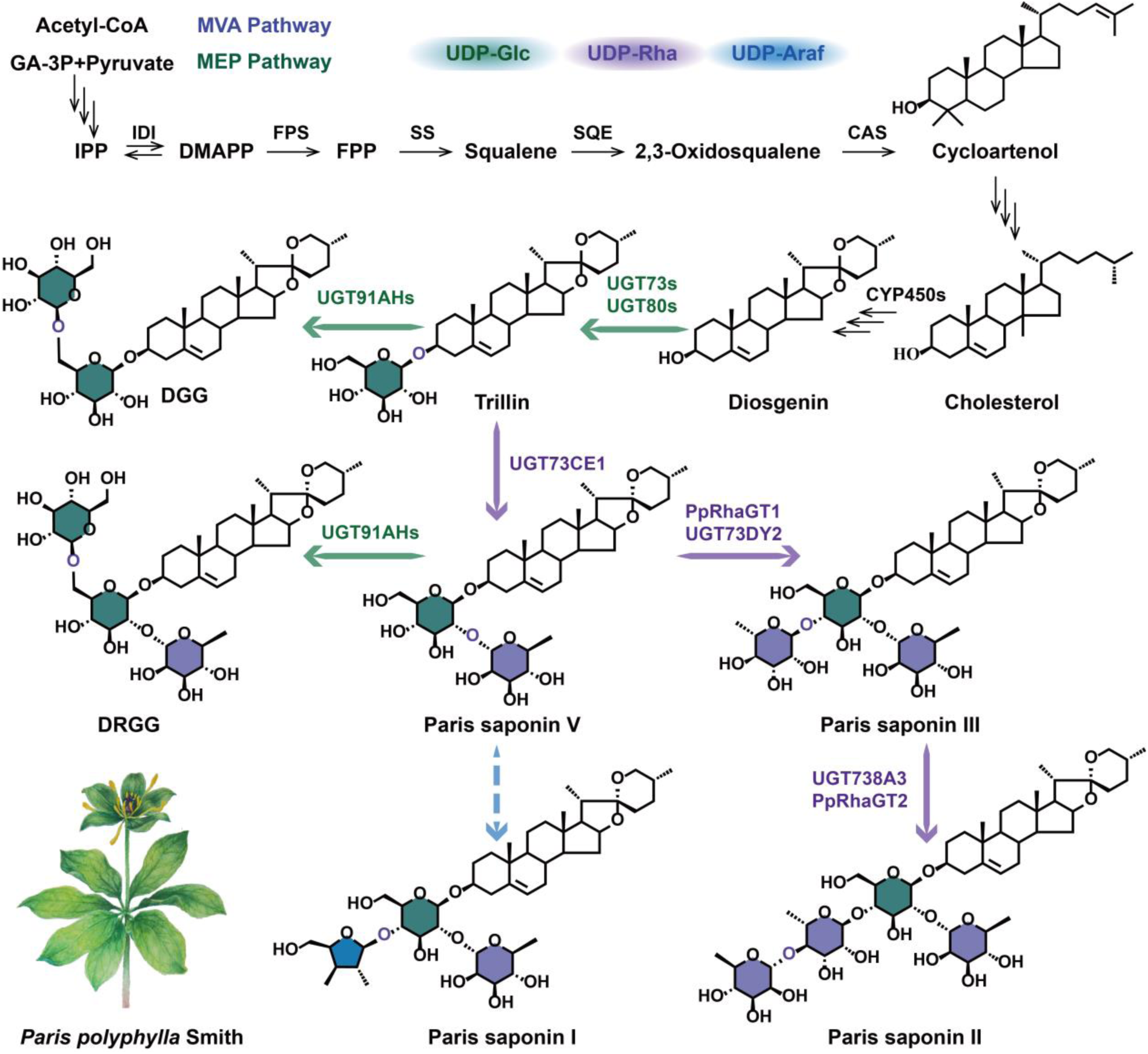
The biosynthetic pathway of the now-analyzed paris saponins. A single arrow denotes a single reaction while multiple arrows represent multi-step catalytic reactions. The dashed arrows represent biological processes that remain to be experimentally characterized. Intermediate product abbreviations: IPP, isopentenyl pyrophosphate; DMAPP, dimethylallyl pyrophosphate; FPP, farnesyl diphosphate.

In recent years, with the development of synthetic biology, heterologous biosynthesis of active compounds in microorganisms has emerged as a promising strategy to address this challenge. However, the downstream glycosylation pathway of paris saponins remains unclear (Figure 1). To date, studies have demonstrated that UGT73 and UGT80 family members catalyze the initial glucosylation at the C3-OH of steroidal aglycones(*10–13*). UGT73CE1, UGT73DY2, *Pp*RhaGT1 and *Pp*RhaGT2 further addition of rhamnose to the corresponding positions leads to the generation of multiple PSs(*12, 14–16*). Our previous study identified UGT738A3 as the terminal rhamnosyltransferase in the paris saponin II(PSII) biosynthetic pathway, thereby enabling the complete elucidation of the tetrasaccharide paris saponin biosynthetic route(*17*). However, all UGTs identified so far that are involved in the biosynthesis of PSs are glucosyltransferases (GlcGT) and rhamnosyltransferases (RhaGT). The functional characterization of arabinofuranosyltransferase (Ara*f*GT) in the biosynthesis of steroidal saponins remains unexplored.

Paris saponin I contains *β*-L-arabinofuranose (Ara*f*), a rare five-membered ring sugar known to significantly enhance the biological activity of natural products. Structural analogs containing Ara*f* are exceedingly rare, being identified in only a handful of triterpene and steroid saponins such as the vaccine adjuvant QS-21(*18*) and ginsenoside Rc(*19*), while arabinopyranose (Ara*p*), though also limited, is documented in a slightly broader range of natural saponins including rosavin(*20*), betulinic acid (*21*), avenacin A-1(*22*), and pseudopasenoside B(*23*). UDP-Ara*f* serves as the direct sugar donor for *O*-linked Ara*f* glycosylation. However, the Ara*f* residue is chemically unstable and susceptible to isomerization to its β-L-arabinopyranose form, making UDP-Ara*f* extremely expensive (approximately $ 519.75 per 0.1 mg) and scarce. Due to the rarity of Ara*f*GT, UGT73CZ2 from *Quillaja saponaria* is the only characterized Ara*f*GT in plants to date(*18*).

To overcome these challenges, we reprogrammed the sugar donor specificity of a steroidal glycoalkaloid rhamnosyltransferase (UGT93M3) through structure-guided engineering to enable Ara*f* transfer activity, and the key amino acid residue (His368) responsible for selective sugar donor conversion was identified. We also integrated UDP-sugar biosynthetic modules(UDP-Rha and UDP-Ara*f*) to address donor availability limitations and reconstructed the paris saponin I biosynthetic pathway in yeast. Subsequently, we compared and integrated multiple functionally characterized UGTs, designed mutants through enzyme engineering approaches to enhance UGT catalytic activity, thereby enabling multi-step glycosylation and the heterologous synthesis of diverse PSs, including diosgenin-3-*O*-glucosyl-(1→6)-glucoside (DGG), diosgenin-3-*O*-rhamnosyl(1→2)[glucosyl(1→6)]glucoside (DRGG), paris saponin I and paris saponin II. This study, for the first time, enabled the successful establishment of the heterologous biosynthetic pathway for paris saponin I. At the mean time, it constructed a microbial platform enabling the *de novo* biosynthesis of a series of PSs, thereby providing a valuable reference for the efficient and scalable production of bioactive steroid saponins.

## 2. Results

### 2.1. Engineering UGT93M3 for efficient arabinofuranosylation of PSI

Recent studies have identified UGT93M3 as a rhamnosyltransferase involved in the solanine biosynthesis pathway of *Solanum nigrum*, which catalyzes the transfer of rhamnose to the C4’-OH of *β*-solamargine, resulting in the formation of *α*-solamargine(*24*) (Figure 2a). Given the structural similarity between *β*-solamargine and paris saponin Ⅴ(PSⅤ) and Ⅵ(PSⅥ), we demonstrated that wild-type UGT93M3 is capable of glycosylating PSⅤ *in vitro*, specifically catalyzing the addition of a rhamnose moiety to the C4’-OH group, resulting in the formation of paris saponin Ⅲ (PSⅢ)(*17*)(Figure 2a). This result confirms that UGT93M3 exhibits a substrate diversity. To further investigate its capacity to utilize other types of sugar donors, we selected a range of sugar donor candidates (e.g., UDP-Glc, UDP-Ara*p*) and performed *in vitro* incubation assays with UGT93M3. When UDP-Ara*p* was used as the glycosyl donor in place of UDP-Rha, an unknown compound was detected under the catalysis of UGT93M3. This compound shared the expected *m*/*z* value ([M+Na]⁺= 877.46) for PSI but displayed distinct chromatographic behavior (Figure 2b). We hypothesized that it is a C4’-arabinopyranosylated derivative, however, structural confirmation was not possible due to the lack of an authentic standard. This observation also suggests that UGT93M3 might possess the potential to utilize UDP-Ara*f* as a glycosyl donor for the catalytic synthesis of PSI.

**Figure 2.**
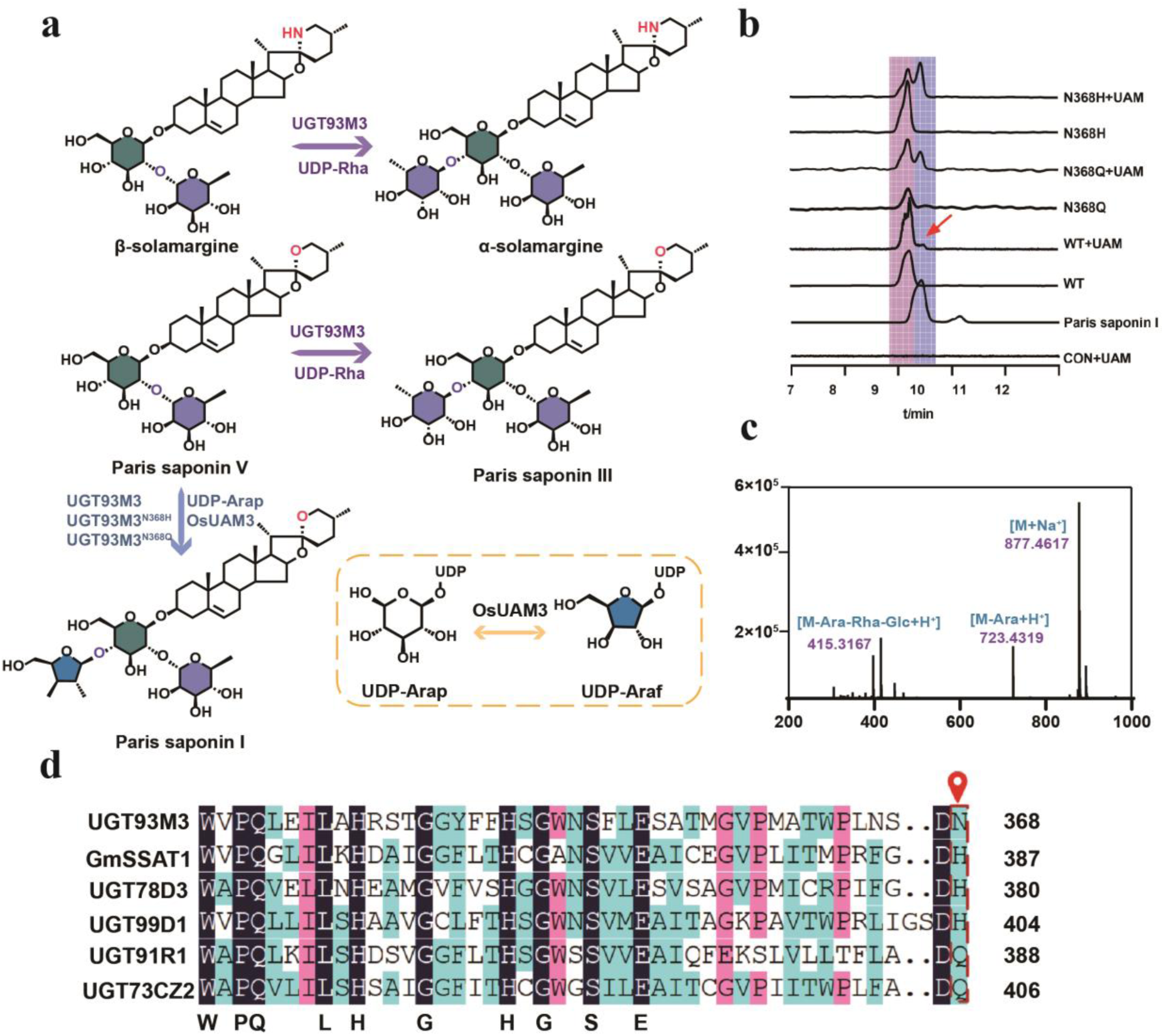
The synthesis of PSⅠ and the enzyme engineering modification of UGT93M3. (a) UGT93M3 utilizes β-solamargine as the substrate and UDP-Rha as the glycosyl donor to catalyze the synthesis of α-solamargine. UGT93M3 utilizes PS Ⅴ as the substrate and UDP-Rha as the glycosyl donor to catalyze the synthesis of PSⅢ. UDP-Ara*p* is converted into UDP-Ara*f* through the catalytic action of UAM. Schematic illustration of the catalytic activity of UGT93M3 variants in the synthesis of PSⅠ. (b) UGT93M3 and its variants were employed in the *in vitro* enzymatic synthesis of PSⅠ. CON: control. UAM: *Os*UAM3 UDP-arabinopyranose mutase 3 (GeneBank ID: NP_001409188.1) WT: wild type of UGT93M3. (c) The characteristic molecular ion peak of PSⅠ. The first-level mass spectrum is [M+Na^+^]. The secondary mass spectra are [M-Ara+H^+^] and [M-Ara-Rha-Glc+H^+^]. (d) Sequence alignment of UGT93M3 with other homologous sequences of AraGTs, Asn^368^ of UGT93M3 exhibits distinct characteristics compared to other AraGTs. UGT93M3: α-solamargine synthase. UGT78D3: *Arabidopsis thaliana* UDP-glucosyl transferase 78D3 (GeneBank ID: NM_121709.2). UGT99D1: *Avena strigosa* UGT99D1 (GeneBank ID: AZQ26921.1). UGT73CZ2: QS-21 biosynthetic Ara*f*GT (GeneBank ID: WLD47571.1). UGT91R1(*31*): (Z)-3-hexenyl β-vicianoside synthase.

PSI (diosgenin-3-*O*-α-L-Rha(l→2)[α-L-Ara(l→4)]-β-D-Glc) contains an arabinofuranose moiety at the C3-OH position of the diosgenin glycan chain. In plants, the biosynthesis of UDP-Ara*f* requires UDP-arabinopyranose mutase (UAM) that catalyzes the transformation of the pyranose ring to form the furanose structure using UDP-Ara*p* as a substrate(*25*) (Figure 2a). Therefore, to investigate the Ara*f*GT activity of UGT93M3, we individually expressed an arabinopyranose mutase (*Os*UAM3 from *Oryza sativa*), which efficiently converts UDP-Ara*p* to UDP-Ara*f*(*26*), along with UGT93M3 in *E. coli*. Enzyme assays were conducted by incubating *Os*UAM3 and UGT93M3 with UDP-Ara*p in vitro*. The cascade allowed *Os*UAM3 to generate UDP-Ara*f* from UDP-Ara*p*, which was then used by UGT93M3. Although only trace amounts of PSI were detected, the result shows that UGT93M3 can use UDP-Ara*f* to glycosylate PSⅤ and forms PSI(Figure 2b,c).

To improve the catalytic efficiency of UGT93M3 in transferring UDP-Ara*f*, we performed site-directed mutagenesis targeting key relevant residues within UGT93M3. Previous studies implicate the C-terminal residue of the plant secondary product glycosyltransferase (PSPG) motif in determining sugar donor specificity(*27–30*). Sequence alignment with known AraGTs revealed that His or Gln residues are conserved at the C-terminus of the PSPG motif, whereas UGT93M3 harbors Asn368 (Figure 2d). Based on this observation, we suspected that replacing the Asn368 with His or Gln would improve UDP-Ara*f* recognition and Ara*f*-transfer capability. To evaluate this hypothesis, site-directed mutagenesis was conducted to generate two UGT93M3 mutants, UGT93M3^N368H^ and UGT93M3^N368Q^. Fortunately, both mutants produced readily detectable PSI when assayed in the *Os*UAM3-coupled system (Figure 2b), demonstrating significantly enhanced Ara*f*-transfer activity compared to the wild-type enzyme.

### 2.2. Demonstration of arabinofuranosyl transfer mechanism based on MD simulation

To further investigate the molecular mechanisms responsible for the enhanced catalytic activity of the mutant, we first determined the enzyme kinetic parameters of both UGT93M3 and the N368H mutant. The *k_ca_*_t_/*K_m_*values clearly demonstrate that the N368H mutant exhibits significantly enhanced catalytic efficiency compared to the wild-type enzyme (Figure S1, Table S1).

Next, molecular docking was employed to elucidate the mechanistic basis for the differences in sugar donor recognition and binding between the wild-type enzyme and the N368H mutant. Molecular docking simulations performed with AlphaFold3 indicated that the shorter side chain of Asn368 in the wild-type enzyme results in a higher binding energy for UDP-Ara*f* (−8.30 kcal·mol^-1^) and places the sugar moiety in a non-productive configuration (Figure 3a). By contrast, the extended imidazole side chain of His in the N368H mutant induced a conformational inversion of the sugar group, substantially reducing the binding energy to −9.47 kcal·mol^-1^ (Figure 3b). Although glutamine at position 368 (N368Q) possesses a side chain of comparable length, it fails to stabilize the catalytically competent pose due to the absence of protonation capability (Figure S2). These findings collectively demonstrate that a histidine residue at the C-terminus of the PSPG motif acts as a critical molecular determinant for AraGT activity. Consistent with this observation, a histidine residue at the equivalent position in *At*UGT78D3 has been previously demonstrated to be essential for both UDP-Ara binding and glycosyltransferase activity(*30, 32*).

**Figure 3.**
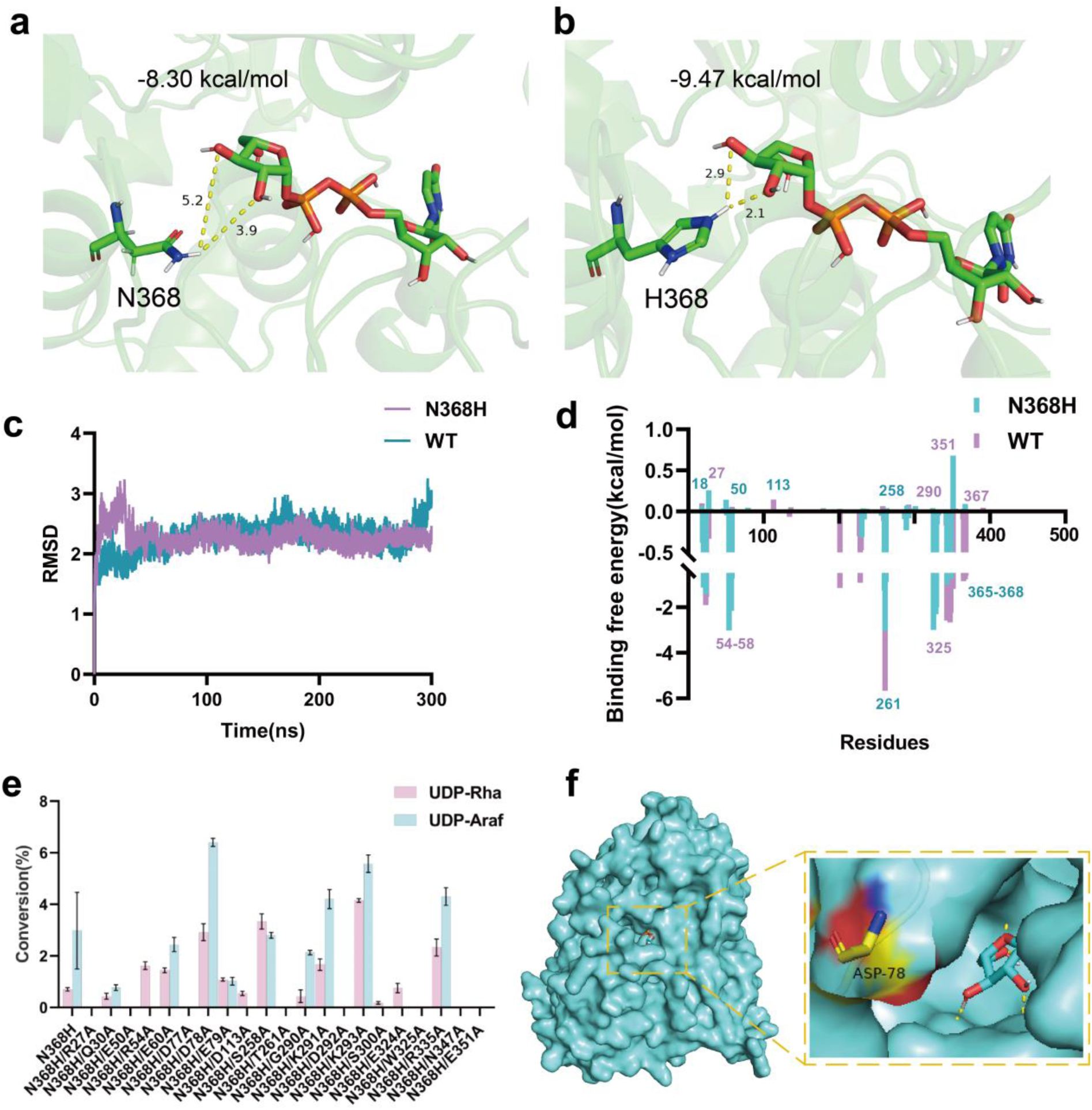
Molecular dynamics simulation and site-directed mutagenesis analysis of the UGT93M3 gene. (a) The docking complex of the WT with UDP-Ara*f*. (b) The docking complex of the N368H with UDP-Ara*f*. The yellow dotted lines represent the hydrogen bonds formed between the protein and the amino acid residues. (c) The RMSD of the wild type and N368H mutant of UGT93M3. (d) The binding free energy of each amino acid at each position in the N368H mutant. (e) The results of alanine scanning mutation detection based on the binding free energy. (f) The Asp78 residue is located at the entrance of the protein’s glycosyl donor channel, and alanine scanning mutations will affect the volume of this channel.

To investigate the role of key amino acids in the dynamic process, we employed 300 ns MD simulations to dissect the dynamic differences between the wild-type enzyme and the N368H variant. The stable trajectory, characterized by minimal RMSD fluctuations, indicates that both simulated systems reached a well-equilibrated state, with the mutant adopting a distinctly more rigid scaffold(Figure 3c), as evidenced by reduced root-mean-square deviation (RMSD) and root-mean-square fluctuation (RMSF) (Figure S3).

Decomposition analysis of free energy contributions revealed distinct differences in UDP-Ara*f* and residue interactions between the wild-type and N368H mutant, leading to the identification of eight key residues (E18, R27, E50, D113, S258, G290, E351, and D367). Residues exhibiting significant changes in energy contributions were predominantly located within the UDP-Ara*f* glycosyl donor binding site. Based on these findings, we thought that rational engineering of residues unfavorable for substrate binding (binding free energy contribution ΔG > 0.003 kcal·mol⁻¹) may enhance enzyme affinity and consequently improve catalytic efficiency. Thus, we performed alanine scanning mutagenesis on the aforementioned residues. In addition, to further validate critical interaction sites, residues T261, W325, and N347, which also exhibited substantial energy contributions, were subjected to alanine scanning. These findings provide evidence that the magnitude of free energy contribution critically influences protein catalysis, as evidenced by the markedly diminished or entirely abolished enzymatic activity in most mutants, such as those at residues R27, E50, D77, T261, D292, W325, and E351. Notably, the D78A variant exhibited a pronounced gain-of-function, enhancing the turnover of both sugar donors. Structural analysis revealed that D78 is positioned at the gateway of the glycosyl donor channel (Figure 3f). Its substitution with alanine likely relieves a potential steric bottleneck, resulting in a modest expansion of the channel aperture that improves glycosyl donor accommodation and boosts catalytic efficiency.

### 2.2. Construction and optimization of the diosgenin biosynthetic pathway in yeast

To evaluate the potential of UGT93M3^N368H^ in microbial cell factories, we attempted to engineer *S. cerevisiae* to construct the heterologous biosynthetic pathway of PSI. The pathway is modularized as two parts: the diosgenin synthesis module and the glycosylation module. In plants, cytochrome P450 monooxygenases (CYP450s) oxidize cholesterol to generate diosgenin(*33–36*). First, starting from the high-yield 2,3-oxidosqualene-producing strain SQ1, we introduced DHCR7 and DHCR24 from *Solanum tuberosum* under the control of the GAL1,10 promoter to redirect metabolic flux toward cholesterol production, yielding strain TZ13, which produced 14.58 mg·L^-1^ cholesterol(Figure 4a,b, Figure S4). Subsequently, the native ERG6 promoter was replaced with the relatively weaker ERG7 promoter to downregulate competing endogenous ergosterol biosynthesis, resulting strain TZ14, enabled significantly increased in cholesterol production to 70.52 mg·L^-1^, which has 3.83-fold increase compared to strain TZ13 (Figure 4b).

**Figure 4.**
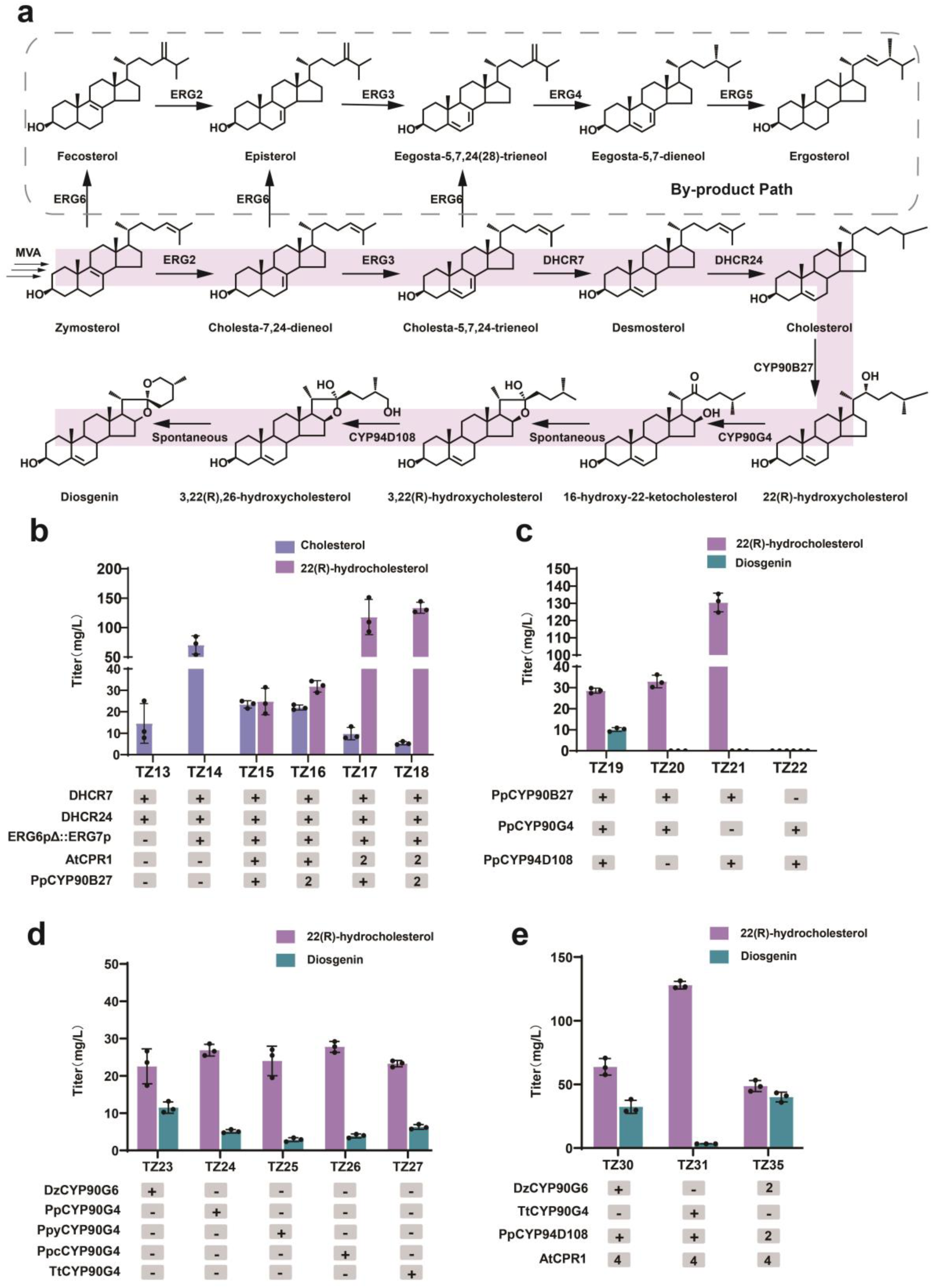
Construction and optimization of the diosgenin biosynthetic pathway in *S. cerevisiae*. (a) The biosynthesis of cholesterol in yeast and the endogenous sterol synthesis pathway in yeast. ERG2: C-8 sterol isomerase. ERG3: C-5 sterol desaturase. ERG4: C-24(28) sterol reductase. ERG5: C-22 sterol desaturase. ERG6: *Δ*(24)-sterol C-methyltransferase. DHCR7: 7-dehydrocholesterol reductase, DHCR24: sterol C-24 reductase, CYP90G4: C16(*S*)-hydroxylase from *P. polyphylla*. CYP94D108: C26-hydroxylase from *P. polyphylla*. (b) Cholesterol was detected in both the TZ13 and TZ14 strains, 22-hydroxycholesterol was detected in TZ15 to TZ18 strains, *At*CPR1: cytochrome P450 reductase from *Arabidopsis thaliana*. CYP90B27: C22(R)-hydroxylase from *P. polyphylla*. (c) Diosgenin was detected exclusively in strains that had simultaneously integrated CYP90B27, CYP90G4, and CYP904108. (d) Functional comparison of *Pp*CYP90G4 Isoenzymes, *Ppy*CYP90G4: C16(S)-hydroxylase from *Ppy*. *Ppc*CYP90G4: C16(S)-hydroxylase from *Ppc*. *Tt*CYP90G4: C16(*S*)-hydroxylase from *Tt*. *Dz*CYP90G6: C16(S)-hydroxylase from *Dz*. (e) *Dz*CYP90G6 and *Pp*CYP94D108 were selected as the enzyme components for the biosynthesis of diosgenin. The yeast strains were incubated for 120 h at 30 ℃, 200 rpm in YPD medium supplemented with 2 % glucose. All values represent the mean ± standard deviation (SD) derived from three biologically independent experiments. The genotypes of all yeast strains constructed in this study can be found in Table S2. The source data underlying figures b-e are provided in Table S3.

Next, we integrated *Pp*CYP90B27 and *At*CPR1 into the strain TZ14 to produce 22(*R*)-hydroxycholesterol, achieving a yield of 24.79 mg·L^-1^. However, 23.45 mg·L^-1^ of cholesterol remained unconverted, indicating that the catalytic capacity was insufficient for complete substrate conversion. To overcome this limitation, we simultaneously overexpressed both genes, resulting in an increased yield to 133.8 mg·L^-1^ (strain TZ18), which has a 4.4-fold increase compared to strain TZ15 (Figure 4b). Diosgenin biosynthesis requires three P450 enzymes, CYP90B27, CYP90G4 and CYP94D108 (Figure S5). Among these, CYP90G4 is a multifunctional enzyme exhibiting both C16(*S*)-hydroxylase and C22(*S*)-hydroxylase activities. Based on these findings, we propose that the coordinated action of CYP90G4 and CYP94D108 may be sufficient to drive the biosynthesis of diosgenin. But systematic combinatorial expression analysis revealed that diosgenin production was strictly dependent on the co-expression of all three P450s in strain TZ19 (Figure 4c), suggesting that 22(*R*)-hydroxycholesterol is an indispensable biosynthetic intermediate. The 22(*S*)-hydroxycholesterol by-product generated during the CYP90G4-catalyzed reaction may lead to substrate waste. To maximize substrate utilization, we screened and compared CYP90G4 orthologs from *Ppy*, *Ppc*,*Tt*, and CYP90G6 from *Dioscorea zingiberensis* (*Dz*) (Figure 4c). Among these, *Dz*CYP90G6 demonstrated superior C16-hydroxylase activity, yielding 11.49 mg·L^-1^ diosgenin compared to 6.27 mg·L^-1^ with *Tt*CYP90G4 (Figure 4d). The codon optimization of *Dz*CYP90G6 and *Tt*CYP90G4 was conducted together with the overexpression of *At*CPR1 in yeast, the resulting strain TZ30 achieved a diosgenin titer of 32.39 mg·L^-1^. Further amplifition of gene copy number in strain TZ35 subsequently improved the titer to 40.10 mg·L^-1^ (Figure 4e). Strain TZ35 exhibited the highest diosgenin synthesis capability and was chosen as the foundational strain for PSI biosynthesis.

### 2.3. Stepwise assembly and optimization of the PSI glycosylation pathway

The glycosylation modules employed in PSI biosynthesis comprise three UGTs: steroid saponin C3-OH GlcGT, C2’-OH RhaGT, and C4’-OH Ara*f*GT. We first identified diosgenin C3-OH GlcGT from *Ppc* that catalyzes the transfer of UDP-Glc to diosgenin, resulting in the formation of trillin. To ensure an adequate supply of substrates for the subsequent glycosylation reaction, we carried out site-directed mutagenesis on *Ppc*UGT4. Structural models generated form AlphaFold 3 were used to guide molecular docking studies (Figure S6). Based on the docking results, alanine-scanning mutagenesis was performed on substrate-proximal residues located within a 5 Å radius of the bound ligands. The catalytic activities of *Ppc*UGT4 mutants were evaluted *in vitro* using diosgenin as substrates, with UDP-Glc serving as the sugar donor during 3-hour reaction periods. Mutations at positions G150, T151, G153 and D154 nearly abolished enzymatic activity, indicating that these residues are essential for catalytic function (Figure S7). Among the single mutants, F211A and L302A exhibited enhanced activity toward diosgenin. Accordingly, we constructed the F211A/L302A double mutant, which displayed even higher catalytic activity. Ultimately, the optimized F211Y/L302T double mutant showed a 1.91-fold increase in activity and achieved 86.5% conversion of diosgenin within 3 hours(Figure S7). The diosgenin-producing stain TZ35 was subsequently engineered with integrating the optimized *Ppc*UGT4^F211Y/L302T^ mutant to enable trillin biosynthesis, resulting in strain TZ36. This mutant-enhanced strain produced 859 μg·L^-1^ trillin, representing a 1.31-fold increase compared to the wild-type *Ppc*UGT4 (371 μg·L^-1^, Figure 5b).

**Figure 5.**
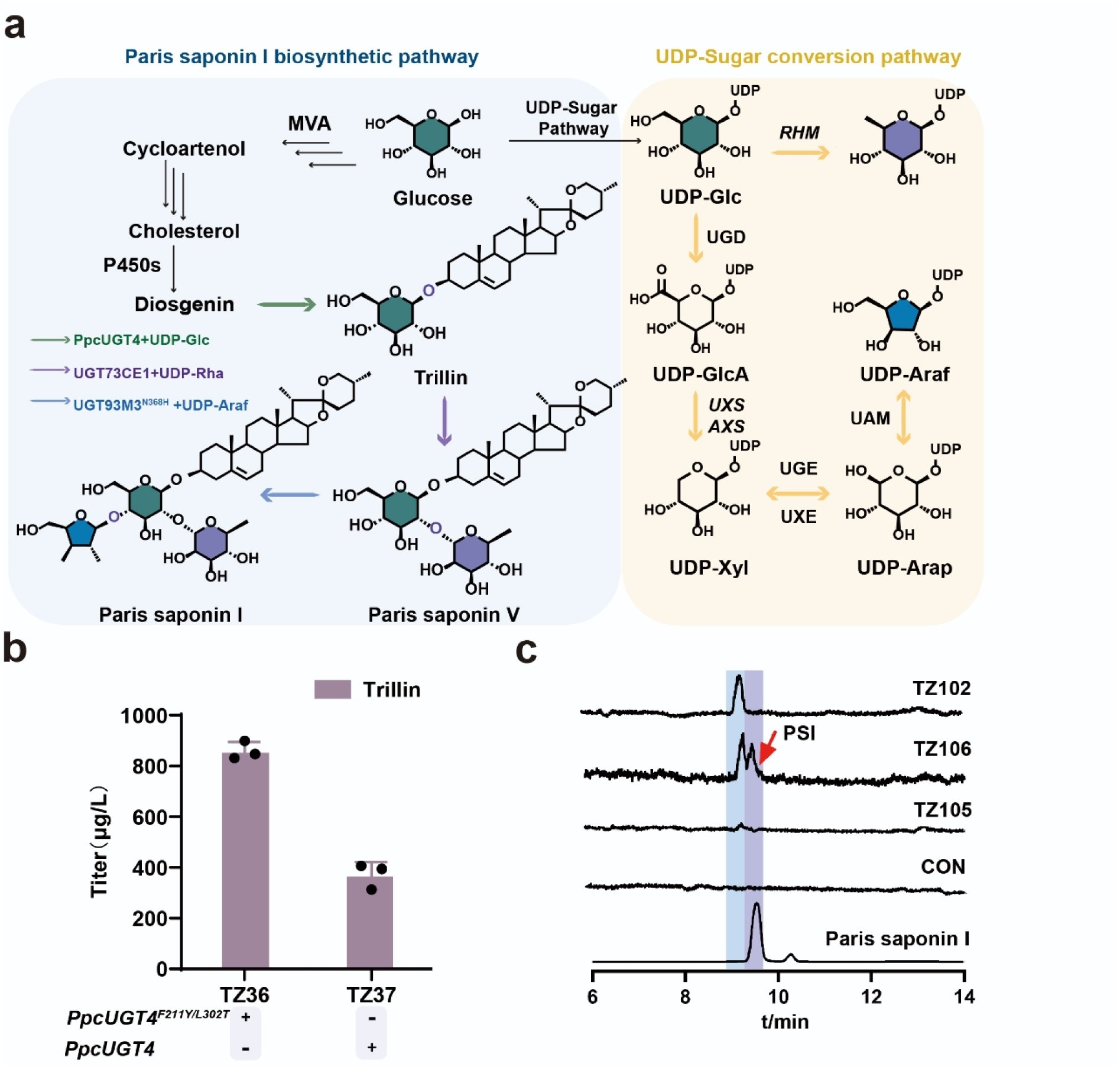
*De novo* biosynthesis of PSⅠ in yeast through integrating the heterologous glycosylation pathway from plants. (a)Schematic representation of PSI synthesis in *S. cerevisiae* and biosynthetic pathways of UDP-Rha and UDP-Ara*f*. This metabolic pathway can be systematically categorized into three distinct phases: diosgenin biosynthesis, UDP-Ara*f* biosynthesis, and PSⅠ biosynthesis. The synthesis of diosgenin is completed under the catalytic action of DHCR (DHCR7 and DHCR24) and P450 (*Pp*CYP90B27, *Dz*CYP90G6 and *Pp*CYP94D108). UDP-Rha is synthesized from UDP-Glc under the catalysis of RHM1. UDP-Ara*f* is synthesized from UDP-Glc under the catalysis of multiple enzymes including UGD, UXS, UXE, UGE and UAM. Finally, PSⅠ is gradually synthesized under the catalysis of *Ppc*UGT4, UGT73CE1 and UGT93M3^N368H^. (b) Strain TZ36 expressing the F211Y/L302T mutant exhibits higher trillin yield than TZ37 harboring the wild-type enzyme. (c) After fermentation, PSⅠ was detected in strain TZ106. The yeast strains were incubated for 120 h at 30 ℃, 200 rpm in YPD medium supplemented with 2 % glucose. All values represent the mean ± SD derived from three biologically independent experiments. The source data underlying figure e is provided in Table S4.

For the biosynthesis of PSⅤ, precursor of PSI, we introduced the UDP-Rha biosynthetic pathway by integrating *Prunus persica* RHM1, which catalyzes the conversion of UDP-Glc to UDP-Rha (Figure 5a)(*37*). UGT73CE1, which transfers rhamnose to the C2’-OH position of trillin to form PSⅤ, was subsequently integrated into strain TZ36, yielding TZ40 that produced 110 μg·L-1 PSⅤ (Figure S8). A functional UDP-Ara*f* supply system is essential for heterologous PSI production in yeast. UDP-Ara*f* is usually synthesized through a multi-enzyme cascade reaction involving four sequential enzymatic steps in plant, catalyzing by UDP-glucose dehydrogenase (UGD), UDP-glucuronic acid decarboxylase (UXS), UDP-xylose 4-epimerase (UXE), and UDP-arabinopyranose mutase (UAM) (Figure 5a). UDP-glucose 4-epimerase (UGE), a cytosolic bifunctional enzyme, facilitates the interconversion between UDP-Xyl and UDP-Ara*p*, as well as between UDP-Glc and UDP-Gal(*38, 39*). To establish a *de novo* UDP-Ara*f* biosynthetic pathway in yeast and identify an optimal UDP-Ara*f* donor-producing strain, we utilized the UGD, UXS, UGE, and UXE from *Arabidopsis* and *Oryza sativa* as query sequences to search for highly homologous genes in *Ppy* and *Tt*. These candidate genes were subsequently cloned and integrated into the PSⅤ producing strain TZ40, generating strains TZ71-74. Building on this foundation, *Os*UAM3 was integrated into the TZ71-74 strains to generate the TZ75-78 strains. Subsequently, due to the high selectivity for UDP-Ara*f*, UGT93M3^N368H^ was introduced into the TZ71-78 strains, resulting in the TZ101-108 strains. After 120 h fermentation in flask, 0.74 μg·L^-1^ PSI was detected in strain TZ106 (Figure 4h). For comparison, PSI was not detected in the TZ102 strain, which lacks *Os*UAM3. In addition, TZ105, which differs from TZ106 solely by the presence of UGE, also failed to produce detectable levels of PSI.

### 2.5. From enzyme mining to strain construction for engineered yeast producing DGG, DRGG and PSII

Although UGT93M3^N368H^ exhibits Ara*f*GT activity and efficiently utilizes UDP-Ara*f*, the native AraGT in *P. polyphylla* remains unidentified. To address this, we used UGT93M3 as a query sequence for phylogenetic analysis of glycosyltransferases in the *P. polyphylla* transcriptome, identifying three candidate UGTs, UGT91AH8-10, that cluster within the same clade as UGT93M3 (Figure S9). However, subsequent cloning and functional characterization revealed that none of these candidates possess Ara*f*GT activity. However, expanded substrate screening uncovered that UGT91AH8-10 catalyzed C6’-OH glucosylation of trillin and PG to yield DGG ([M+Na]^+^ *m/z* 761.3782) and PGG ([M+Na]^+^ *m/z* 777.406). When PSⅤ and PSⅥ were used as acceptors, UGT91AH10 produced DRGG ([M+Na]^+^ *m/z* 907.4304) and PRGG ([M+Na]^+^ *m/z* 923.4325). Structures were confirmed by NMR of purified DGG, comparison with a DRGG standard, and mass spectral matching to literature data(Figure S10-S19, Table S5-S6).

Previous studies reported that DGG exhibits notable hemolytic activity, whereas DRGG displays potent antifungal properties(*12, 40*). To enable efficient synthesis of these two active compounds in yeast, we performed enzyme engineering on UGT91AH10. The three-dimensional structure of UGT91AH10 was modeled with AlphaFold 3 (Figure S20) and used to guide mutagenesis. Enzyme activity was assayed with PSⅤ or trillin as acceptor and UDP-Glc as sugar donor. Alanine scanning identified seven residues (H20, D119, Y120, G151, E202, E284, H375) essential for catalysis, as their mutation abolished activity. Sequence alignment with homologs was then performed to guide efficiency improvements(*12, 41*), following which fourteen residues were selected for site-directed mutagenesis. The results showed that only the I13L and V117L significantly improved catalytic efficiency *in vitro*. When trillin was used as the substrate, the I13L exhibited a 1.84-fold increase in activity compared to the wild-type enzyme, whereas the V117L showed a 3.78-fold enhancement using PSⅤ as the substrate (Figure S21).

These variants (I13L for DGG biosynthesis; V117L for DRGG biosynthesis) were integrated as key enzymatic components in engineered yeast strains (Figure 6a). Then UGT91AH10, UGT91AH10^I13L^, and UGT91AH10^V117L^ were separately integrated into the trillin producing strain TZ36 and the PSV producing strain TZ40, resulting in the engineered strains TZ60-65 (Figures 6c). After shake-flask fermentation, the strain TZ61 exhibited the highest DGG production at 150 μg·L^-1^, while strain TZ65 achieved the maximum DRGG titer of 85 μg·L^-1^.

**Figure 6.**
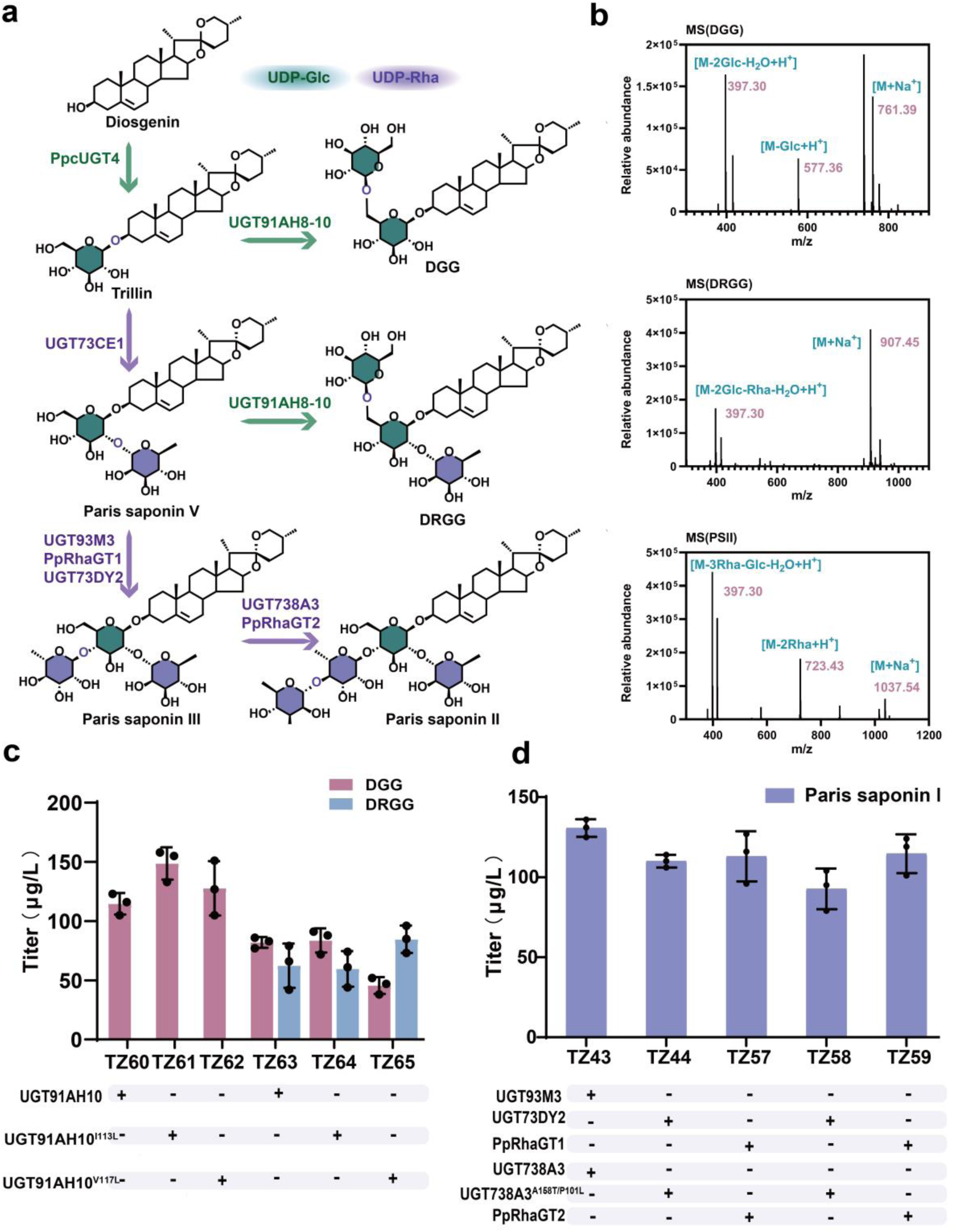
Construction of yeast strains for production of DGG, DRGG and PSⅡ. (a) Schematic representation of DGG, DRGG and PSⅡ synthesis in *S. cerevisiae*. The biosynthesis of DGG, DRGG and PSⅡ exclusively utilizes UDP-Glc and UDP-Rha as sugar donors. DGG and DRGG are synthesized from trillin and PSⅤ, respectively, through the catalytic activity of UGT91AH10. PSⅢ is synthesized from PSⅤ, through the catalytic activity of UGT93M3, UGT73DY2 and *Pp*RhaGT1. PSⅡ is synthesized from PSⅢ, through the catalytic activity of UGT738A3 and *Pp*RhaGT2. (b) The characteristic molecular ion peaks of DGG, DRGG and PSⅡ. (d) The integrated I13L mutant exhibits the highest DGG yield. The integrated V117L mutant exhibits the highest DRGG yield. (e) Comparison of the activities of C4’-OH and C4′-OH rhamnosyltransferases related to PSⅡ synthesis. The TZ43 strain, integrating UGT93M3 and UGT738A3^P101L/A158T^, exhibits the highest PSⅡ yield. The yeast strains were incubated for 120 h at 30 ℃, 200 rpm in YPD medium supplemented with 2 % glucose. All values represent the mean ± SD derived from three biologically independent experiments. The source data underlying figures b-e are provided in Table S7.

Furthermore, PSII is an active steroid saponin with a tetrasaccharide chain at the C3-OH position in *P. polyphylla*, which suppresses colorectal carcinogenesis by regulating mitochondrial fission and the NF-*κ*B pathway(*42*). Nowadays, the biosynthetic pathway of PSII have been elucidated(*15, 16*). To establish heterologous biosynthesis of PSII in yeast, we integrated the PSⅢ biosynthetic genes UGT93M3, *Pp*RhaGT1, and UGT73DY2(*15, 16, 24*), together with the PSII biosynthetic genes UGT738A3 and *Pp*RhaGT2(*16, 43*), into the PSⅤ-producing strain TZ40, resulting in the generation of a series of PSII-producing strains (Figure 6d). Comparative analysis revealed that UGT93M3 demonstrated superior catalytic efficiency compared to UGT73DY2 and *Pp*RhaGT1 in PSⅢ synthesis, while the UGT738A3^P101L/A158T^ mutant(*17*) (UGT738A3^P101L/A158T^ is an enzymatic variant that exhibits enhanced catalytic activity compared to the WT enzyme previously characterized in our laboratory) outperformed *Pp*RhaGT2 in the final glycosylation step. The optimal strain TZ43 achieved a PSII titer of 130 μg·L^-1^, representing the highest yield among all engineered strains producing PSII (Figure 6d).

## 3. Discussion

In this study, we successfully improved the sugar donor selectivity of the RhaGT UGT93M3, conferring Ara*f*GT activity by enzyme engineering. To date, the endogenous AraGT in *P. polyphylla* remains unidentified. Our findings provide a valuable reference for its future discovery and functional characterization, and offer a transferable strategy for constructing biosynthetic pathways of other natural active compounds containing arabinofuranosyl moieties. Furthermore, four PSs biosynthetic pathways were reconstructed in engineered yeast, leading to the first heterologous synthesis of PSII (130 μg·L^-1^), PSI (0.74 μg·L^-1^), DGG (150 μg·L^-1^), and DRGG (85 μg·L^-1^), thus laying the foundation for the sustainable and scalable production of steroidal saponins.

Ara*f* is a vital component of polysaccharides in bacteria, protozoa, fungi, and plants (e.g., constituting 5-10% of cell wall polysaccharides in *A. thaliana* and *O. sativa*) (*44*), yet it is rarely found in natural products. In plants, UAM catalyzes the reversible ring contraction of UDP-Ara*p* to form UDP-Ara*f*(*45*). However, the isomerization rate from UDP-Ara*f* to UDP-Ara*p* proceeds approximately ten times faster than the reverse reaction. Consequently, UDP-Ara*p* to UDP-Ara*f* reach a molar ratio of approximately 9:1 at thermodynamic equilibrium, which limits the accumulation of the furanose-type donor(*46–49*). This inherent instability likely contributes to the low PSI yield observed in our engineered yeast system. Furthermore, the *de novo* synthesis of UDP-Ara*f* is metabolically demanding, requiring four enzymatic steps starting from UDP-Glc. Crowe *et al.* expressed seven genes encoding the salvage pathway in *S. cerevisiae* to directly activate free sugars by exogenous supplementation of arabinose, enabling the biosynthesis of UDP-Ara*p* and UDP-Ara*f*. However, UDP-Ara*f* accumulates only at very low levels (∼0.037 μmol·g^-1^ DCW) (*38*). To address this challenge, we cloned the UGD, UXS, UGE, and UXE genes from two closely related plant species (*Ppy* and *Tt*), established four UDP-Ara*p* biosynthetic pathway strains (TZ101, TZ102, TZ103, TZ104) through combinatorial assembly of enzyme modules, and introduced the functionally characterized *Os*UAM3 gene. Ultimately, PSI was detected in one of these strains (TZ106), which contained enzyme components derived from *Tt*UXS1, *Tt*UGD3, and *Tt*UXE2. The key distinction between TZ105 and TZ106 resides in the presence of UXE in TZ106 as opposed to UGE in TZ105. In contrast, UXE is predominantly localized in the Golgi apparatus in plant cells and specifically catalyzes the interconversion between UDP-Xyl and UDP-Ara*p*. In our study, the strain expressing UXE achieved detectable levels of PSI accumulation. These findings suggest that both the functional characteristics of enzyme components and the subcellular environment are critical determinants in metabolic engineering. Therefore our work establishes a valuable framework for the heterologous reconstruction of polysaccharide glycosylation steps (particularly those involving Ara*f* moieties) within complex natural product biosynthetic pathways.

Currently, several studies are focused on reconstructing the biosynthetic pathways of steroidal saponins in plant chassis systems. For example, Studies have successfully reconstructed the biosynthetic pathways of PSⅢ and PSII in *Nicotiana benthamiana*, resulting in a maximum PSs yield of 93.64 DCW(*15, 50*). However, yeast chassis systems offer distinct advantages, including facile genetic manipulation, rapid growth, well-established fermentation processes, and low production costs, highlighting their potential for the heterologous production of bioactive compounds(*51, 52*). In previous studies on the heterologous production of steroid compounds, *S. cerevisiae* has been engineered for the synthesis of steroidal precursors (e.g., cholesterol, diosgenin)(*53–55*), as well as certein steroidal hormone(e.g., progesterone, ergosterol et al.)(*56, 57*). But the PSs targeted in this study hold different structure from these steroidal compounds, which the PSs normally have the features with a characteristic spiroketal side chain at the C-17 position (e.g., pennogenin, diosgenin) and the extensive glycosylation specifically at the C-3 hydroxyl group(*58*). This essential C3 glycosylation, absent in precursors and hormones. This behavior not only enhances the water solubility and bioavailability of steroid compounds, but also determines the diverse pharmacological activities of PSs, which can be attributed to the structural diversity of sugar chains in steroid saponins, including variations in sugar types and quantities (e.g., UDP-Glc, UDP-Rha, and UDP-Ara)(*58*).

To the best of our knowledge, this study reports the first microbial synthesis of highly glycosylated steroidal saponins in yeast, achieving titers of 130 μg·L^-1^ PSII, 0.74 μg·L^-1^ PSI, 150 μg·L^-1^ DGG, and 85 μg·L^-1^ DRGG. Through the identification of enzymes and subsequent enzyme engineering modifications, we have successfully integrated 16 genes into yeast, enabling the synthesis of heterologous compounds within a relatively long biosynthetic pathway. However, the current yield of PSs remains low. To optimize the sugar donor pathway and enhance the yield of the final product, we deleted the endogenous yeast GAL7 gene to redirect metabolic flux. Additionally, the PGM2 gene was overexpressed to increase the availability of UDP-Glc precursors (Figure S22). However, when these modifications were introduced into the PSII-producing strain TZ43, the PSII yield unexpectedly decreased to 50 µg·L^-1^ (Figure S23). This reduction may result from the disruption of metabolic homeostasis within yeast cells, which can lead to detrimental effects. To address such metabolic imbalances in future metabolic engineering endeavors, potential strategies include precise modulation of gene expression levels (e.g., through promoter engineering or copy number adjustment), aiming to achieve a more balanced metabolic flux distribution(*59*).

In summary, this study presents a successful example of the discovery and functional characterization of the rare Ara*f*GTs and its application in engineered biosynthetic pathways, which provides a valuable reference for the subsequent identification and characterization of specific UGTs. Furthermore, this study highlights yeast as a potential production platform, marking a significant advance for the efficient synthesis of a wide range of steroid saponins.

## 4. Methods

### 4.1. Plant material

*Paris polyphylla* var*. yunnanensis* specimens were collected from Yongsheng County, Lijiang City, Yunnan Province, China.

*Paris polyphylla* var*. chinensis* specimens were collected from Huangshan City, Anhui Province, China.

*Trillium. tschonoskii* plant material was collected in April 2023 from Baoxing County, Ya’an City, Sichuan Province, China.

Following collection, all samples were immediately flash-frozen in liquid nitrogen and subsequently stored at −80 °C. Three independent biological replicates were prepared for each sample for subsequent sequencing.

### 4.2. UPLC-Q-TOF/MS and UPLC-Q-Exactive HF/MS analysis

UPLC-Q-TOF/MS Method: Fermentation samples were analyzed using a UPLC-Q-TOF/MS system (Waters SYNAPT G2-Si). Chromatographic separation was performed on a C18 reversed-phase column (Waters; 1.7 μm, 2.1 mm × 100 mm) maintained at 40 °C, with an injection volume was 3 μL. The detection was carried out using a diode array detector (DAD). The mobile phase consisted of water (A) and acetonitrile (B), delivered at a flow rate of 0.3 mL/min. Gradient elution was employed as follows: 0–2 min, 50–70% B; 2–7 min, 70–80% B; 7–10 min, 80–95% B; 10–16 min, 95–95% B; 16–18 min, 95–100% B; 18–20 min, 100% B.

Mass spectrometric detection was performed using the Xevo Q-TOF mass spectrometer operated in positive ion electrospray ionization (ESI+) mode. Full-scan mass spectra were acquired over the m/z range 50 to 550. The capillary voltage was set to 1 kV, and the cone voltage was 40 V. Nitrogen was used as the nebulization and desolvation gas. The source temperature was 120 °C, the desolvation gas temperature was 450 °C, the desolvation gas flow was 800 L/h, and the cone gas flow was 50 L/h. Data acquisition and analysis were controlled by MassLynx 4.1 software to obtain accurate mass information.

UPLC-Q-Exactive HF/MS Method: Chromatographic separation used an identical gradient program and column to the UPLC-Q-TOF/MS method described above. The MS spectrometric analysis was performed on a Q-Exactive HF mass spectrometer operated in positive electrospray ionization(ESU+) mode under the following parameters: full scan range m/z 100–1000 in positive ion mode; MS¹ resolution 120,000 with automatic gain control (AGC) target 3e6; MS² resolution 30,000 with AGC target 1e5; sheath gas flow 45 arbitrary units (arb); auxiliary gas flow 15 arb; spray voltage 3000 V; capillary temperature 320°C; ion lens RF level (S-Lens) 50%; and ion source temperature 350 °C.

### 4.3. Protein Purification and Enzyme Kinetic Parameter Determination

The target UGT and its mutant proteins, expressed as His-MBP fusions from the pET28a vector in *E. coli* Transetta(DE3) cells were purified. Large-scale culture were induced with IPTG (1 mM), and crude enzyme extracts were prepared from the harvested cells.

Recombinant proteins were purified by immobilized metal affinity chromatography (IMAC) using Ni-NTA resin. The detailed purification protocol was as follows: The clarified crude protein extract was loaded onto a Ni-NTA column pre-equilibrated with binding buffer (50 mM Tris-HCl, pH=7.4). Bound proteins were eluted stepwise using equilibration buffer containing increasing imidazole concentrations (10 mM, 20 mM, 50 mM, 100 mM, 200 mM, 300 mM, and 500 mM). Eluates fractions from each step were collected separately. Fractions containing the target protein, identified by SDS-PAGE analysis, were pooled and concentrated to approximately 1 mL using a 30 kDa molecular weight cut-off (MWCO) centrifugal ultrafiltration device. The concentrated protein was aliquoted, flash-flozen in liquid nitrogen, and stored at −80°C for subsequent kinetic assays.

Enzyme kinetic parameters ( *K*_m_) for the acceptor substrate were determined a concentration gradient (1.25, 2.5, 5, 10, 20, 40, 80, 100, 200, 300, and 500 μM). Reactions were performed in triplicate in a total volume of 50 μL, containing 5 μg of purified enzyme, 100 mM UDP-glucose (UDP-Glc), the specified concentration acceptor substrate, and 50 mM Tris-HCl buffer (pH7.4). After incubation for 1 h at 30°C, reactions were terminated by adding 150 μL of ice-cold methanol. The mixture was centrifuged at 15,000 rpm for 30 min, and the resulting supernatant was analyzed by UPLC-Q-Exactive HF/MS as described previously. Kinetic parameters (*V*_max_ and *K*_m_) were derived by non-linear regression analysis of the data using GraphPad Prism software (version 10).

### 4.4. Homology Modeling, Docking Analysis, and Site-Directed Mutagenesis

The three-dimensional structure of the target glycosyltransferase was generated by homology modeling using AlphaFold 3 (http://alphafoldsever.com/). Ligand structures (acceptor substrate) and the sugar donor (UDP-glucose) were retrieved from the PubChem database(https://pubchem.ncbi.nlm.nih.gov/compound). Molecular docking simulations were performed using AutoDock Vina, with the homology model as the receptor and the ligands as substrates. Optimal binding poses were selected based on docking scores and subjected to structural analysis and visualization using PyMOL. To investigate functional residues, alanine scanning mutagenesis was performed on all residues within 5 Å of the bound substrate in the optimal docing pose, followed by iterative mutagenesis of key positions identified in initial screens. Site-directed mutations were introduced into the His-MBP-pET28a-target glycosyltransferase plasmid using a Fast Mutagenesis System kit. Verified mutant plasmids were transformed into *E.coli Transetta* (DE3) competent cells, with preliminary mutant viability assessed through transformation efficiency assays prior to protein expression and functional characterization. The mutation primers used in this study can all be found in Table S8, Table S9 and Table S10.

### 4.5. *In vitro* biosynthesis of PSI

UGT93M3, its mutant variants, and OsUAM3 proteins were expressed as His-MBP fusion proteins using the pET28a vector in Escherichia coli Transetta(DE3) cells. Large-scale cultures were induced with IPTG (1 mM), and enzyme extracts were subsequently prepared from the harvested cells. PSI biosynthesis reactions contained 50 mM Tris-HCl buffer (pH 7.4), 5 mM MgCl₂, 1 mM UDP-Arap, and 0.2 mM PSⅤ in a total volume of 50μL. Reactions were initiated by 23μL of crude enzyme extract (UGT93M3 or its mutants) and incubated at 30°C. After 1 hour, 24 μL of crude *Os*UAM3 enzyme was added. The reaction proceeded for an additional 9 h (10 h total duration). The reaction was terminated by adding 150 μL of ice-cold methanol. The mixture was centrifuged at 15000 rpm for 30 min, and the resulting supernatant was analyzed by UPLC-Q-Exactive HF/MS as described previously.

### 4.6. Construction of recombinant strains

The genotypes of all yeast strains constructed in this study can be found in Table S1. Except for genes from Figure 3d, all genes employed in this study for yeast strain construction were codon-optimized, and their corresponding sequences are provided in Supplementary Data 2. All the genes to be synthesized and the primers required for cloning were provided by Beijing Tianyi Huiyuan Co., Ltd. The yeast genomic loci and homologous recombination fragments utilized in this study are detailed in Table S11 and Supplementary Data 3.

Pick single yeast colonies from YPD solid plates and inoculate them into 2 mL of YPD liquid medium. Incubate at 30 °C with shaking for 16 hours. Transfer the entire volume of the culture to 20 mL of synthetic dropout liquid medium and continue incubation at 30 °C with shaking for 5–6 hours (A600 nm approximately 0.8–2.0). Under a laminar flow hood, transfer the culture to a 50 mL centrifuge tube and centrifuge at 3600 rpm at room temperature for 5 minutes to pellet the cells. Discard the supernatant, resuspend the cells in 10 mL of sterile water, centrifuge again, and repeat this washing step twice. Next, resuspend the cells in 600 µL of sterile water, transfer to 1.5 mL microcentrifuge tubes, and centrifuge at 4 °C and 3600 rpm for 3 min. Discard the supernatant and collect the cell pellet. Resuspend the cells in 500 µL of 1 M lithium acetate (LiAc), aliquot into 1.5 mL tubes at 50 µL per tube, and centrifuge again at 4 °C and 3600 r·min⁻¹ for 3 minutes. Discard the supernatant to obtain competent cells. The transformation mixture contains 50% PEG3350, 1 M LiAc, denatured ssDNA (heated at 100°C for 5 min and immediately cooled on ice for 2 min), target DNA, and ddH₂O. All transformation steps are performed on ice. If not used immediately, competent cells can be resuspended in 100 µL of 20% sterile glycerol and stored at −80 °C.

After adding the transformation mixture, incubate the samples at 30°C for 30 minutes, followed by heat shock at 42°C for 15 min. Centrifuge at 3600 rpm for 5 min, discard the supernatant, resuspend the cells in 100 µL of sterile water, and spread onto SD-Ura dropout plates. After culturing for 2–3 days, pick PCR-confirmed positive colonies and transfer them to 5-FOA plates to eliminate the plasmid, yielding the final recombinant strains.

### 4.7. Fermentation of Recombinant Strains and Sample Preparation

Positive recombinant clones were picked from YPD solid plates (2% glucose) and used to inoculate 2 mL of YPD liquid medium (2% glucose). After 16h incubation at 30 °C with shaking (200 rpm), the optical density (OD_600_) was measured. A portion of the culture was then transferred to 20 mL of fresh YPD medium (2% glucose) and fermented for 5 days at 30 °C with 200 rpm orbital shaking.

Cells were harvested by centrifugation ( 4000 rpm, 5 min), and the cell pellet was resuspended in 5 mL of 20% (w/v) potassium hydroxide in 50%(v/v) aqueous ethanol. Cell lysis was achieved by boiling the suspension for 10 min. The cooled lysate was extracted three times with an equal volume of ethyl acetate. After centrifugation, the combined organic supernatants were dehydrated using anhydrous sodium sulfate. The extract was concentrated to dryness under a gentle nitrogen stream at ≤30°C. The residue was reconstituted in 200 μL of methanol, filtered throuth a 0.22-μm membrane, transferred to a GC vial, and analyzed by LC-MS analysis for metabolite profiling.

## Supplemental inpormation

Supplemental information is available at bioRxiv Online.

Document Supplemental inpormation. Figures S1–S23, and Tables S1–S11.

Supplementary Data 1. The nucleotide sequences of the genes identified in this study.

Supplementary Data 2. The codon-optimized sequence used in this study.

Supplementary Data 3. The upstream and downstream homologous repair fragments of the genomic loci used in this study.

## Data availability statement

Data supporting the findings of this study are available within the article and its Supplementary Information files.

## Acknowledgements

We thank the Core Facilities Center of Capital Medical University for supporting sample testing in this research.

## Author contributions

Haowen Wang, Yuxin Yang and Ziya Wu contributed equally to this work. Haowen Wang, Xianan Zhang, Yating Hu, Yuxin Yang and Ziya Wu conceived the project and wrote the manuscript. Haowen Wang, Xianan Zhang, Yating Hu, Yuxin Yang, Ziya Wu, Chi Zhang, Chenxing Sun, Zihan Yu, Bowen Qiu, Xuan Liu designed and performed all the experiments. Haowen Wang, Yuxin Yang, Ziya Wu, Huan Zhao, Yinying Ba, Chi Zhang, Chenxing Sun, Zihan Yu, Bowen Qiu analyzed the results.

