## Supplementary information for "Elucidation and engineering of arabinofuranosyltransferase to enable total *de novo* biosynthesis of paris saponins in yeast"

Chi Zhang

Peking University Shougang Hospital, Beijing 100000, China

Haowen Wang, Yuxin Yang and Ziya Wu contributed equally to this work.

^*^Corresponding Authors

(Xianan Zhang).

(Yating Hu).

**Supplementary Tables**

Table S 1 The enzyme kinetic parameters of UGT93M3 and N368H

| Gene | Substrate | *K*_m_(μM) | *V*_max_(nM/min) | *K*_cat_(s^−1^) | R^2^ |
| --- | --- | --- | --- | --- | --- |
| UGT93M3 | Polyphyllin Ⅴ | 108.7 | 0.62 | 0.03 | 0.9501 |
| UGT93M3 | UDP-Ara*f* | 0.23 | 0.71 | 0.01 | 0.7610 |
| UGT93M3 | UDP-Rha | 0.04 | 103.3 | 9.57 | 0.7472 |
| UGT93M3^N368H^ | Polyphyllin Ⅴ | 231.5 | 2.6 | 0.05 | 0.8248 |
| UGT93M3^N368H^ | UDP-Ara*f* | 0.67 | 9.57 | 0.59 | 0.7863 |
| UGT93M3^N368H^ | UDP-Rha | 0.64 | 5.4 | 0.33 | 0.9003 |

Table S 2 The genotypes of all yeast strains constructed in this study

| **Strain** | **Parent strain** | **Genotype** | **Reference** |
| --- | --- | --- | --- |
| CEN.PK113-5D | -- | MATa ura3-52 trp1-289 leu2-3, 112 his3Δ1 MAL2‑8c SUC2 | Retained in the laboratory |
| SQ1 | CEN.PK113-5D | CAN1Δ:: pTEF1-CAS9-tCYC1, Gal80Δ, X3:: pTDH3-ERG20-tCYC1-pPGK1-ERG9-tFBAl, BTS1Δ:: pTHD3-tHMG1-tFBA1-pTEF1-ERG1-tIDP1 | Retained in the laboratory |
| TZ13 | SQ1 | XI3:: tCYC1-DHCR7-pGAL10,pGAL1-DHCR24-tADH1 | This study |
| TZ14 | TZ13 | ERG6Δ:: ERG7pr-ERG6 | This study |
| TZ15 | TZ14 | V1:: tPGK1-PpCYP90B27-pGAL10,pGAL1-AtCPR1-tADH1 | This study |
| TZ16 | TZ14 | III1:: tPGK1-PpCYP90B27-pGAL10,pGAL1 | This study |
| TZ17 | TZ14 | III1:: pGAL10,pGAL1-AtCPR1-tADH1 | This study |
| TZ18 | TZ14 | III1:: tPGK1-PpCYP90B27-pGAL10,pGAL1-AtCPR1-tADH1 | This study |
| TZ19 | TZ18 | V3:: tTEF1-CYP90G4-pGAL10,pGAL1-CYP94D108-tHXT7 | This study |
| TZ20 | TZ18 | V3:: tTEF1-CYP90G4-pGAL10,pGAL1 | This study |
| TZ21 | TZ18 | V3:: pGAL10,pGAL1-CYP94D108-tHXT7 | This study |
| TZ22 | TZ14 | V3:: tTEF1-CYP90G4-pGAL10,pGAL1-CYP94D108-tHXT7 | This study |
| TZ23 | TZ18 | V3:: tTEF1-DzCYP90G6-pGAL10,pGAL1-CYP94D108-tHXT7 | This study |
| TZ24 | TZ18 | V3:: tTEF1-PpCYP90G4-pGAL10,pGAL1-CYP94D108-tHXT7 | This study |
| TZ25 | TZ18 | V3:: tTEF1-PpyCYP90G4-pGAL10,pGAL1-CYP94D108-tHXT7 | This study |
| TZ26 | TZ18 | V3:: tTEF1-PpcCYP90G4-pGAL10,pGAL1-CYP94D108-tHXT7 | This study |
| TZ27 | TZ18 | V3:: tTEF1-TtCYP90G4-pGAL10,pGAL1-CYP94D108-tHXT7 | This study |
| TZ30 | TZ18 | XII2:: tADH1-AtCPR1-pGAL10,pGAL1-AtCPR1-tADH1, V3:: tTEF1-DzCYP90G6-pGAL10,pGAL1-CYP94D108-tHXT7 | This study |
| TZ31 | TZ18 | XII2:: tADH1-AtCPR1-pGAL10,pGAL1-AtCPR1-tADH1, V3:: tTEF1-TtCYP90G4-pGAL10,pGAL1-CYP94D108-tHXT7 | This study |
| TZ35 | TZ30 | XI5:: tTEF1-DzCYP90G6-pGAL10,pGAL1-CYP94D108-tHXT7 | This study |
| TZ36 | TZ35 | XI2:: tADH1-PpcUGT4^F211Y/L302T^-pGAL10,pGAL1 | This study |
| TZ37 | TZ35 | XI2:: tADH1-PpcUGT4-pGAL10,pGAL1 | This study |
| TZ40 | TZ36 | XII1:: tTEF1-UGT73CE1-pGAL10,pGAL1-RHM1-tADH1 | This study |
| TZ71 | TZ40 | XII4:: tTEF1-TtUXS1-pGAL10,pGAL1-TtUGD3-tHXT7 | This study |
| TZ72 | TZ40 | XII4:: tTEF1-TtUXS1-pGAL10,pGAL1-TtUGD3-tHXT7 | This study |
| TZ73 | TZ40 | XII4:: tTEF1-PpUXS3-pGAL10,pGAL1-PpUGD1-tHXT7 | This study |
| TZ74 | TZ40 | XII4:: TEF1-TtUXS1-pGAL10,pGAL1-TtUGD3-tHXT7 | This study |
| TZ75 | TZ71 | X4:: pGAL10,pGAL1-TtUGE1-tHXT7 | This study |
| TZ76 | TZ72 | X4:: pGAL10,pGAL1-TtUXE2-tHXT7 | This study |
| TZ77 | TZ73 | X4:: pGAL10,pGAL1-PpUGE2-tHXT7 | This study |
| TZ78 | TZ74 | X4:: pGAL10,pGAL1-PpUXE1-tHXT7 | This study |
| TZ101 | TZ75 | XII2:: tTEF1-UGT93M3^N368H^-pGAL10,pGAL1 | This study |
| TZ102 | TZ76 | XII2:: tTEF1-UGT93M3^N368H^-pGAL10,pGAL1 | This study |
| TZ103 | TZ77 | XII2:: tTEF1-UGT93M3^N368H^-pGAL10,pGAL1 | This study |
| TZ104 | TZ78 | XII2:: tTEF1-UGT93M3^N368H^-pGAL10,pGAL1 | This study |
| TZ105 | TZ75 | XII2:: tTEF1-UGT93M3^N368H^-pGAL10,pGAL1-OsUAM3-tHXT7 | This study |
| TZ106 | TZ76 | XII2:: tTEF1-UGT93M3^N368H^-pGAL10,pGAL1-OsUAM3-tHXT7 | This study |
| TZ107 | TZ77 | XII2:: tTEF1-UGT93M3^N368H^-pGAL10,pGAL1-OsUAM3-tHXT7 | This study |
| TZ108 | TZ78 | XII2:: tTEF1-UGT93M3^N368H^-pGAL10,pGAL1-OsUAM3-tHXT7 | This study |
| TZ43 | TZ40 | XII4:: tTEF1-UGT93M3-pGAL10,pGAL1-UGT738A3^P101L/A158T^-tADH1 | This study |
| TZ44 | TZ40 | XII4:: tTEF1-UGT93M3-pGAL10,pGAL1-UGT738A3-tADH1 | This study |
| TZ57 | TZ40 | XII4:: tTEF1-PpRhaGT1-pGAL10,pGAL1-UGT738A3^P101L/A158T^-tADH1 | This study |
| TZ58 | TZ40 | XII4:: tTEF1-PpRhaGT1-pGAL10,pGAL1-PpRhaGT2-tADH1 | This study |
| TZ59 | TZ40 | XII4:: tTEF1-UGT73DY2-pGAL10,pGAL1-UGT738A3^P101L/A158T^-tADH1 | This study |
| TZ60 | TZ36 | XII4:: pGAL10,pGAL1-UGT91AH10-tADH1 | This study |
| TZ61 | TZ36 | XII4:: pGAL10,pGAL1-UGT91AH10^I13L^-tADH1 | This study |
| TZ62 | TZ36 | XII4:: pGAL10,pGAL1-UGT91AH10^V117L^-tADH1 | This study |
| TZ63 | TZ40 | XII4:: pGAL10,pGAL1-UGT91AH10-tADH1 | This study |
| TZ64 | TZ40 | XII4:: pGAL10,pGAL1-UGT91AH10^I13L^-tADH1 | This study |
| TZ65 | TZ40 | XII4:: pGAL10,pGAL1-UGT91AH10^V117L^-tADH1 | This study |

Table S 3 The data of the yeast strains in Figure 4

| Strain name | Cholesterol(mg/L) | | | 22(R)-hydrocholesterol(mg/L) | | | Diosgenin(mg/L) | | | OD_600_ | | |
| --- | --- | --- | --- | --- | --- | --- | --- | --- | --- | --- | --- | --- |
| TZ13 | 10.78 | 25.11 | 7.85 | N.D. | N.D. | N.D. | N.D. | N.D. | N.D. | 10.13 | 10.59 | 10.98 |
| TZ14 | 72.81 | 84.83 | 53.91 | N.D. | N.D. | N.D. | N.D. | N.D. | N.D. | 9.97 | 10.67 | 10.34 |
| TZ15 | 21.56 | 23.81 | 24.97 | 23.81 | 31.39 | 19.17 | N.D. | N.D. | N.D. | 11.06 | 10.56 | 10.47 |
| TZ16 | 21.69 | 21.02 | 23.26 | 28.94 | 34.28 | 32.27 | N.D. | N.D. | N.D. | 10.84 | 10.85 | 11.69 |
| TZ17 | 8.42 | 7.88 | 13.09 | 109.58 | 93.29 | 151.31 | N.D. | N.D. | N.D. | 10.45 | 10.33 | 9.36 |
| TZ18 | 5.12 | 4.76 | 6.23 | 130.05 | 126.82 | 144.55 | N.D. | N.D. | N.D. | 10.27 | 9.89 | 10.58 |
| TZ19 | N.D. | N.D. | N.D. | 28.59 | 27.36 | 29.79 | 10.88 | 10.38 | 9.10 | 10.24 | 9.68 | 11.03 |
| TZ20 | N.D. | N.D. | N.D. | 36.17 | 30.23 | 32.42 | N.D. | N.D. | N.D. | 10.87 | 10.56 | 10.21 |
| TZ21 | N.D. | N.D. | N.D. | 131.29 | 124.67 | 135.40 | N.D. | N.D. | N.D. | 9.47 | 10.05 | 10.45 |
| TZ22 | N.D. | N.D. | N.D. | N.D. | N.D. | N.D. | N.D. | N.D. | N.D. | 10.13 | 10.18 | 10.65 |
| TZ23 | N.D. | N.D. | N.D. | 23.58 | 26.61 | 17.43 | 11.03 | 10.26 | 13.18 | 10.38 | 9.57 | 10.85 |
| TZ24 | N.D. | N.D. | N.D. | 26.60 | 28.58 | 25.47 | 5.03 | 5.63 | 4.75 | 9.22 | 11.00 | 9.74 |
| TZ25 | N.D. | N.D. | N.D. | 19.51 | 27.01 | 25.50 | 2.35 | 2.92 | 3.40 | 10.13 | 10.09 | 9.49 |
| TZ26 | N.D. | N.D. | N.D. | 26.29 | 27.83 | 29.21 | 4.01 | 3.33 | 4.35 | 10.33 | 9.85 | 10.69 |
| TZ27 | N.D. | N.D. | N.D. | 22.35 | 23.42 | 24.02 | 6.16 | 6.99 | 5.74 | 9.04 | 10.42 | 9.93 |
| TZ30 | N.D. | N.D. | N.D. | 70.66 | 63.18 | 57.64 | 29.09 | 38.30 | 29.79 | 9.32 | 10.69 | 10.55 |
| TZ31 | N.D. | N.D. | N.D. | 131.32 | 125.92 | 126.61 | 3.29 | 3.35 | 3.38 | 10.34 | 10.36 | 9.97 |
| TZ35 | N.D. | N.D. | N.D. | 53.27 | 48.37 | 44.57 | 35.90 | 41.06 | 43.34 | 10.47 | 10.23 | 10.11 |

Table S 4 The data of the yeast strains in Figure 5

| Strain name | Trillin(μg/L) | | | OD_600_ | | |
| --- | --- | --- | --- | --- | --- | --- |
| TZ36 | 831.35 | 899.25 | 847.65 | 10.39 | 9.98 | 11.02 |
| TZ37 | 313.75 | 406.59 | 394.18 | 10.64 | 10.57 | 10.43 |

Table S 5 Enzyme kinetic parameters of UGT91AH8/9/10 with trillin as substrate

| UGT | *K*_m_(μM) | *V*_max_(μM·min^-1^) | *k*_cat_(s^-1^) | *k*_cat_/*K*_m_ |
| --- | --- | --- | --- | --- |
| UGT91AH8 | 5.97 | 0.030 | 5.01 | 0.84 |
| UIGT91AH9 | 6.94 | 0.017 | 2.83 | 0.41 |
| UGT91AH10 | 9.65 | 0.012 | 1.99 | 0.21 |

Table S 6 Enzyme kinetic parameters of UGT91AH8/9/10 with paris saponin V as substrate

| UGT | *K*_m_(μM) | *V*_max_(μM·min^-1^) | *k*_cat_(s^-1^) | *k*_cat_/*K*_m_ |
| --- | --- | --- | --- | --- |
| UGT91AH8 | 8.34 | 0.022 | 3.67 | 0.44 |
| UIGT91AH9 | 6.29 | 0.020 | 3.33 | 0.53 |
| UGT91AH10 | 14.34 | 0.013 | 2.17 | 0.15 |

Table S 7 The data of the yeast strains in Figure 6

| Strain name | DGG(μg/L) | | | DRGG(μg/L) | | | Paris saponin Ⅱ(μg/L) | | | OD_600_ | | |
| --- | --- | --- | --- | --- | --- | --- | --- | --- | --- | --- | --- | --- |
| TZ43 | N.D. | N.D. | N.D. | N.D. | N.D. | N.D. | 136.52 | 125.32 | 131.22 | 10.48 | 10.79 | 10.58 |
| TZ44 | N.D. | N.D. | N.D. | N.D. | N.D. | N.D. | 110.62 | 106.25 | 114.98 | 10.23 | 9.87 | 9.56 |
| TZ57 | N.D. | N.D. | N.D. | N.D. | N.D. | N.D. | 96.22 | 127.13 | 116.98 | 10.58 | 10.56 | 10.03 |
| TZ58 | N.D. | N.D. | N.D. | N.D. | N.D. | N.D. | 79.32 | 104.05 | 95.22 | 9.93 | 10.91 | 10.47 |
| TZ59 | N.D. | N.D. | N.D. | N.D. | N.D. | N.D. | 124.23 | 119.87 | 101.26 | 9.76 | 9.88 | 10.44 |
| TZ60 | 116.01 | 123.22 | 105.02 | N.D. | N.D. | N.D. | N.D. | N.D. | N.D. | 10.58 | 10.23 | 10.62 |
| TZ61 | 133.53 | 158.26 | 155.56 | N.D. | N.D. | N.D. | N.D. | N.D. | N.D. | 10.21 | 10.39 | 10.77 |
| TZ62 | 105.98 | 151.98 | 127.75 | N.D. | N.D. | N.D. | N.D. | N.D. | N.D. | 9.99 | 10.24 | 10.28 |
| TZ63 | 86.61 | 77.65 | 83.45 | 42.56 | 66.98 | 79.03 | N.D. | N.D. | N.D. | 10.57 | 10.56 | 10.34 |
| TZ64 | 88.84 | 91.32 | 72.04 | 61.95 | 74.87 | 44.45 | N.D. | N.D. | N.D. | 10.91 | 10.69 | 10.78 |
| TZ65 | 52.99 | 47.87 | 38.11 | 85.03 | 73.65 | 96.42 | N.D. | N.D. | N.D. | 10.11 | 9.82 | 10.29 |

Table S 8 Primers required for the UGT93M3 mutation experiment

| Name | Sequence (5’-3’) |
| --- | --- |
| R27A-F | caaatccctcttctcgctctctctcaa |
| R27A-R | gcgagaagagggatttggtgtccatat |
| Q30A-F | cttctccgtctctctgcactcatctca |
| Q30A-R | gcagagagacggagaagagggatttgg |
| E50A-F | gtgcacgaaatcatgcactaaaacaa |
| E50A-R | gcatgatttcgtgcaccgatgctcat |
| E60A-F | aacaaaatcaacaagCattccccaac |
| E60A-R | Gcttgttgattttgttgacgttgttt |
| D77A-F | cgtcaaaggccaacgCcgatgaagag |
| D77A-R | Gcgttggcctttgacgttattgaaag |
| D78A-F | caaaggccaacgacgCtgaagaggat |
| D78A-R | Gcgtcgttggcctttgacgttattga |
| E79A-F | aggccaacgacgatgcagaggattac |
| E79A-R | gcatcgtcgttggcctttgacgttat |
| D113A-F | tagtcattatccatgCcgctatgatg |
| D113A-R | Gcatggataatgactactcttttatc |
| S258A-F | tcagttatattcgttGcatttggaac |
| S258A-R | Caacgaatataactgagttaattgat |
| N263A-F | tcatttggaactgcaGCcagtttctca |
| N263A-R | GCtgcagttccaaatgaaacgaatata |
| G290A-F | tctgggtgctaagagCaaaagataaa |
| G290A-R | Gctcttagcacccagatgaatttttg |
| K291A-F | tgggtgctaagaggaGCagataaaaag |
| K291A-R | GCtcctcttagcacccagatgaatttt |
| D292A-F | tgctaagaggaaaagCtaaaaaggga |
| D292A-R | Gcttttcctcttagcacccagatgaa |
| K293A-F | ctaagaggaaaagatGCaaagggagaa |
| K293A-R | GCatcttttcctcttagcacccagatg |
| S300A-F | ggagaaaatcttacaGCtgaaatacag |
| S300A-R | GCtgtaagattttctccctttttatct |
| E324A-F | gaatcatcgctagagCatgggtacct |
| E324A-R | Gctctagcgatgattcctctatgatt |
| R335A-F | gagattctagcacatGCttctactggt |
| R335A-R | GCatgtgctagaatctctagttgaggt |
| N347A-F | ttccattcgggatggGCttccttccta |
| N347A-R | GCccatcccgaatggaaaaaataacca |

Table S 9 Primers required for the *Ppc*UGT4 mutation experiment

| Name | Sequence (5’-3’) |
| --- | --- |
| D154A-F | GTTGGTACACGTGGAGCTGTTCAACCTT |
| D154A-R | GCTCCACGTGTACCAACAATAAGAATAA |
| F204-A | GCTTCAGCCAGAACTTTTGGGTCTCCTC |
| F204A-F | AAAGTTCTGGCTGAAGCCATGGTCAAGA |
| F211A-F | GTCAAGAATAAAGGGGCCTTACCTTCTT |
| F211A-R | GCCCCTTTATTCTTGACCATGAATTCAG |
| F277A-F | CCGATACACATATTCGCCACAATGCCAT |
| F277A-R | GCGAATATGTGTATCGGAACCTTTAGCG |
| F489A-F | ACTATCGTACCTTTCGCTGGAGATCAAC |
| F489A-R | GCGAAAGGTACGATAGTAGTTGGACATG |
| G150A-F | GTTATTCTTATTGTTGCTACACGTGGAG |
| G150A-R | GCAACAATAAGAATAACTATTTGTATAG |
| G153A-F | ATTGTTGGTACACGTGCAGATGTTCAAC |
| G153A-R | GCACGTGTACCAACAATAAGAATAACTA |
| L201S-F | GGAGACCCAAAAGTTTCGGCTGAATTCA |
| L201S-R | GAAACTTTTGGGTCTCCTCCTAAGGGGT |
| L291A-F | GAATTTCCACATCCTGCCTCTCGTGTCA |
| L291A-R | GCAGGATGTGGAAATTCACTAGTTGGCG |
| L302A-F | CCAGCTGGATATAGAGCTTCTTACCAAA |
| L302A-R | GCTCTATATCCAGCTGGCTGCTTGACAC |
| M205A-F | GTTCTGGCTGAATTCGCGGTCAAGAATA |
| M205A-R | GCGAATTCAGCCAGAACTTTTGGGTCTC |
| M226A-F | ATTCAACGAAAGCAAGCGAAGGAAATCA |
| M226A-R | GCTTGCTTTCGTTGAATAGCAATTTCTG |
| M279A-F | CACATATTCTTCACAGCGCCATGGACGC |
| M279A-R | GCTGTGAAGAATATGTGTATCGGAACCT |
| P198A-F | CCCCTTAGGAGGAGACGCAAAAGTTCTG |
| P198A-R | CGTCTCCTCCTAAGGGGTAAAATTCCAG |
| P258A-F | CATTATTGCGAATCCCGCGGCATATGGG |
| P258A-R | CGGGATTCGCAATAATGGCATCTGCTTT |
| Q222A-F | TCAGAAATTGCTATTGCACGAAAGCAAA |
| Q222A-R | GCAATAGCAATTTCTGAAGGTGAAGAAG |
| T151A-F | TATTCTTATTGTTGGTGCACGTGGAGAT |
| T151A-R | CACCAACAATAAGAATAACTATTTGTAT |
| A302C-F | CAGCTGGATATAGATGTTCTTACCAAAT |
| A302C-R | CATCTATATCCAGCTGGCTGCTTGACAC |
| A302D-F | AGCTGGATATAGAGATTCTTACCAAATT |
| A302D-R | TCTCTATATCCAGCTGGCTGCTTGAC |
| A302F-F | CAGCTGGATATAGATTTTCTTACCAAAT |
| A302F-R | AATCTATATCCAGCTGGCTGCTTGACAC |
| A302G-F | AGCTGGATATAGAGGTTCTTACCAAATT |
| A302G-R | CCTCTATATCCAGCTGGCTGCTTGAC |
| A302H-F | CAGCTGGATATAGACATTCTTACCAAAT |
| A302H-R | TGTCTATATCCAGCTGGCTGCTTGACAC |
| A302M-F | GCCAGCTGGATATAGAATGTCTTACCAAATTG |
| A302M-R | CATTCTATATCCAGCTGGCTGCTTGACACG |
| A302S-F | CAGCTGGATATAGATCTTCTTACCAAAT |
| A302S-R | ATCTATATCCAGCTGGCTGCTTGAC |
| A302T-F | CAGCTGGATATAGAACTTCTTACCAAAT |
| A302T-R | TTCTATATCCAGCTGGCTGCTTGAC |
| A302V-F | AGCTGGATATAGAGTTTCTTACCAAATT |
| A302V-R | ACTCTATATCCAGCTGGCTGCTTGAC |
| A302W-F | GCCAGCTGGATATAGATGGTCTTACCAAATTG |
| A302W-R | CCATCTATATCCAGCTGGCTGCTTGACACG |
| A211C-F | GTCAAGAATAAAGGGTGCTTACCTTCTT |
| A211C-R | CACCCTTTATTCTTGACCATGAATTCAG |
| A211D-F | GTCAAGAATAAAGGGGACTTACCTTCTT |
| A211D-R | TCCCCTTTATTCTTGACCATGAATTCAG |
| A211G-F | GTCAAGAATAAAGGGGGCTTACCTTCTT |
| A211G-R | CCCCCTTTATTCTTGACCATGAATTCAG |
| A211H-F | GTCAAGAATAAAGGGCACTTACCTTCTT |
| A211H-R | TGCCCTTTATTCTTGACCATGAATTCAG |
| A211I-F | GTCAAGAATAAAGGGATCTTACCTTCTT |
| A211I-R | ATCCCTTTATTCTTGACCATGAATTCAG |
| A211K-F | GTCAAGAATAAAGGGTGGTTACCTTCTT |
| A211K-R | CTTCCCTTTATTCTTGACCATGAATTCAG |
| A211L-F | GTCAAGAATAAAGGGCTCTTACCTTCTT |
| A211L-R | AGCCCTTTATTCTTGACCATGAATTCAG |
| A211M-F | GTCAAGAATAAAGGGATGTTACCTTCTT |
| A211M-R | CATCCCTTTATTCTTGACCATGAATTCAG |
| A211N-F | GTCAAGAATAAAGGGAACTTACCTTCTT |
| A211N-R | TTCCCTTTATTCTTGACCATGAATTCAG |
| A211P-F | GTCAAGAATAAAGGGCCCTTACCTTCTT |
| A211P-R | GCCCTTTATTCTTGACCATGAATTCAG |
| A211Q-F | GTCAAGAATAAAGGGCAGTTACCTTCTT |
| A211Q-R | CTGCCCTTTATTCTTGACCATGAATTCAG |
| A211R-F | GTCAAGAATAAAGGGCGCTTACCTTCTT |
| A211R-R | CGCCCTTTATTCTTGACCATGAATTCAG |
| A211S-F | GTCAAGAATAAAGGGTCCTTACCTTCTT |
| A211S-R | ACCCTTTATTCTTGACCATGAATTCAG |
| A211V-F | GTCAAGAATAAAGGGGTCTTACCTTCTT |
| A211V-R | ACCCCTTTATTCTTGACCATGAATTCAG |
| A211W-F | GTCAAGAATAAAGGGTGGTTACCTTCTT |
| A211W-R | CCACCCTTTATTCTTGACCATGAATTCAG |
| A211Y-F | GTCAAGAATAAAGGGTACTTACCTTCTT |
| A211Y-R | TACCCTTTATTCTTGACCATGAATTCAG |
| A302C-F | CAGCTGGATATAGATGTTCTTACCAAAT |
| A302C-R | CATCTATATCCAGCTGGCTGCTTGACAC |
| A302D-F | AGCTGGATATAGAGATTCTTACCAAATT |
| A302D-R | TCTCTATATCCAGCTGGCTGCTTGAC |
| A302F-F | CAGCTGGATATAGATTTTCTTACCAAAT |
| A302F-R | AATCTATATCCAGCTGGCTGCTTGACAC |
| A302G-F | AGCTGGATATAGAGGTTCTTACCAAATT |
| A302G-R | CCTCTATATCCAGCTGGCTGCTTGAC |
| A302H-F | CAGCTGGATATAGACATTCTTACCAAAT |
| A302H-R | TGTCTATATCCAGCTGGCTGCTTGACAC |
| A302M-F | GCCAGCTGGATATAGAATGTCTTACCAAATTG |
| A302M-R | CATTCTATATCCAGCTGGCTGCTTGACACG |
| A302S-F | CAGCTGGATATAGATCTTCTTACCAAAT |
| A302S-R | ATCTATATCCAGCTGGCTGCTTGAC |
| A302T-F | CAGCTGGATATAGAACTTCTTACCAAAT |
| A302T-R | TTCTATATCCAGCTGGCTGCTTGAC |
| A302V-F | AGCTGGATATAGAGTTTCTTACCAAATT |
| A302V-R | ACTCTATATCCAGCTGGCTGCTTGAC |
| A302W-F | GCCAGCTGGATATAGATGGTCTTACCAAATTG |
| A302W-R | CCATCTATATCCAGCTGGCTGCTTGACACG |

Table S 10 Primers required for the UGT91AH10 mutation experiment

| Name | Sequence (5’-3’) |
| --- | --- |
| D119A-F | GATTGGCTCGTCTACGCCTACGCCGCCT |
| D119A-R | GCGTAGACGAGCCAATCGGGATTCTCG |
| D154A-F | TCTTCGGTCCGTTGGCCAAACCCCCAAG |
| D154A-R | GCCAACGGACCGAAGAAGCTGAGGGTGG |
| E202A-F | GCATCCGGAATGTCTGCATCCCTTCGGC |
| E202A-R | GCAGACATTCCGGATGCGGGCTTCGGGT |
| E284A-F | GTTGCATTTGGGAGCGCGGCGAAACTAT |
| E376A-F | ATTGCCGATGGTGCATGCGCAAGGGCTC |
| E376A-R | GCATGCACCATCGGCAATACCACCATTG |
| F142A-F | TGCCTACCTCAGCCTAGCCAGCGCCGCC |
| F142A-R | GCTAGGCTGAGGTAGGCAAAGGGGACG |
| F150A-F | CCACCCTCAGCTTCGCCGGTCCGTTGG |
| F150A-R | GCGAAGCTGAGGGTGGCGGCGCTGAAT |
| G19A-F | CCATGGCTTGCCATGGCCCACATCCTCC |
| G19A-R | GCCATGGCAAGCCATGGGATCACCAC |
| G151A-F | ACCCTCAGCTTCTTCGCTCCGTTGGAC |
| G151A-R | GCGAAGAAGCTGAGGGTGGCGGCGCTG |
| H20A-F | TGGCTTGCCATGGGCGCCATCCTCCCCT |
| H20A-R | GCGCCCATGGCAAGCCATGGGATCACC |
| H375A-F | GTATTGCCGATGGTGGCTGAGCAAGGGC |
| H375A-R | GCCACCATCGGCAATACCACCATTGGC |
| K155A-F | TTCGGTCCGTTGGACGCACCCCCAAGAT |
| K155A-R | GCGTCCAACGGACCGAAGAAGCTGAGGG |
| L153A-F | AGCTTCTTCGGTCCGGCGGACAAACCCC |
| L153A-R | GCCGGACCGAAGAAGCTGAGGGTGGCG |
| L159A-F | GGACAAACCCCCAAGAGCGTCCCCCGAG |
| L159A-R | GCTCTTGGGGGTTTGTCCAACGGACCG |
| L187A-F | TCGAGGCTCAGGCCGCGTTGAAGCAAAG |
| L187A-R | GCGGCCTGAGCCTCGAAGAGCCTGAAGG |
| L206A-F | CTGAATCCCTTCGGGCTGCGAATGGAT |
| L206A-R | GCCCGAAGGGATTCAGACATTCCGGATG |
| L206F-F | GTCTGAATCCCTTCGGTTTGCGAATGGAT |
| L206F-R | ACCGAAGGGATTCAGACATTCCGGATGC |
| P152A-F | CTCAGCTTCTTCGGTGCGTTGGACAAAC |
| P152A-R | CACCGAAGAAGCTGAGGGTGGCGGCG |
| P156A-F | CGGTCCGTTGGACAAAGCCCCAAGATTG |
| P156A-R | CTTTGTCCAACGGACCGAAGAAGCTGAG |
| P157A-F | TCCGTTGGACAAACCCGCAAGATTGTCC |
| P157A-R | CGGGTTTGTCCAACGGACCGAAGAAGCT |
| R94A-F | CACCGCCACTACCTGGCGATCGCCTTCG |
| R94A-R | GCCAGGTAGTGGCGGTGGAGATCGTCG |
| R158A-F | TTGGACAAACCCCCAGCATTGTCCCCCG |
| R158A-R | GCTGGGGGTTTGTCCAACGGACCGAAGA |
| R211A-F | CTTGCGAATGGATTCGCAAAGTGCCCTT |
| R211A-R | GCGAATCCATTCGCAAGCCGAAGGGATT |
| R284A-R | GCGCTCCCAAATGCAACATAAACTACAG |
| S191A-F | GCCCTGTTGAAGCAAGCCCTCGACCCG |
| S191A-R | GCTTGCTTCAACAGGGCCTGAGCCTCG |
| S203A-F | ATCCGGAATGTCTGAAGCCCTTCGGCTT |
| S203A-R | CTTCAGACATTCCGGATGCGGGCTTCG |
| S203F-F | TCCGGAATGTCTGAATTCCTTCGGCTTG |
| S203F-R | AATTCAGACATTCCGGATGCGGGCTTCG |
| W15A-F | TGTTGTGGTGATCCCAGCGCTTGCCATGG |
| W15A-R | GCTGGGATCACCACAACATGAAGGCTTC |
| Y120A-F | TGGCTCGTCTACGACGCCGCCGCCTACT |
| Y120A-R | GCGTCGTAGACGAGCCAATCGGGATTCT |
| C70R-F | CGAATTCTCTTTGTCCCGCATCGAACAC |
| C70R-R | GGGACAAAGAGAATTCGACTAGAGTTAG |
| H375Y-F | GGTATTGCCGATGGTGTATGAGCAAGGG |
| H375Y-R | ACACCATCGGCAATACCACCATTGGCAG |
| I13L-F | CCTTCATGTTGTGGTGCTCCCATGGCTT |
| I13L-R | GCACCACAACATGAAGGCTTCCATGGC |
| L164F-F | TGTCCCCCGAGGAGTTCACCAATGTCCC |
| L164F-R | CAACTCCTCGGGGGACAATCTTGGGGG |
| L206F-F | GTCTGAATCCCTTCGGTTTGCGAATGG |
| L206F-R | ACCGAAGGGATTCAGACATTCCGGATGC |
| L414V-F | CAAGTGTTTGAGGTTGGTGATGGTGGAT |
| L414V-R | CCAACCTCAAACACTTGGCGATCCCCTC |
| P254S-F | CCGACTCCGGTCCTGTCCGAAGGAAGT |
| P254S-R | ACAGGACCGGAGTCGGAGGGGGGAAGAG |
| Q76E-F | CATCGAACACCTTCCAGAATCCATCGAG |
| Q76E-R | CTGGAAGGTGTTCGATGCAGGACAAAG |
| Q101E-F | GCCTTCGACTCTTTCGAGGTCCAACTC |
| Q101E-R | CGAAAGAGTCGAAGGCGATCCGCAGGT |
| R237G-F | ATTAAACGAGTTGTATGGAAAACCGGTC |
| R237G-R | CATACAACTCGTTTAATAGCTCTATCC |
| T250A-F | ACTCTTCCCCCCTCCGGCTCCGGTCCTG |
| T250A-R | CCGGAGGGGGGAAGAGTCCGATGGGGAT |
| V117L-F | GAATCCCGATTGGCTCCTCTACGACTAC |
| V117L-R | GGAGCCAATCGGGATTCTCGCGACGGAG |
| V240I-F | GTTGTATCGAAAACCGATCATCCCCATC |
| V240I-R | TCGGTTTTCGATACAACTCGTTTAATAG |
| Y430C-F | GCCAAAGCTAGGGAGTGTAAGCAAGTGC |
| Y430C-R | CACTCCCTAGCTTTGGCTCTGATCTTC |

Table S 11 The genomic loci utilized in this study, along with their corresponding gRNA N20 sequences

| Genomic loci | N20 sequence |
| --- | --- |
| III1 | TTAGCAACGCTCAGGGACTC |
| V1 | CGTTTATAGACGGCACTGTC |
| V3 | CGTCTAGAAGAACAGCACAT |
| X4 | CGCCATTCAAGAGCAGCAAC |
| XI2 | GCTTTACTTGTGGAAGTTCA |
| XI3 | ATATGTCTCTAATTTTGGAA |
| XI5 | TGAGAATACTGTTGTAAAAC |
| XII1 | GGTATGTGCAGTTGATTCAC |
| XII2 | TGAAACTCTAATCCTACTAT |
| XII4 | CTTTATGCATAGAGCTAATT |

**Supplementary Figure**

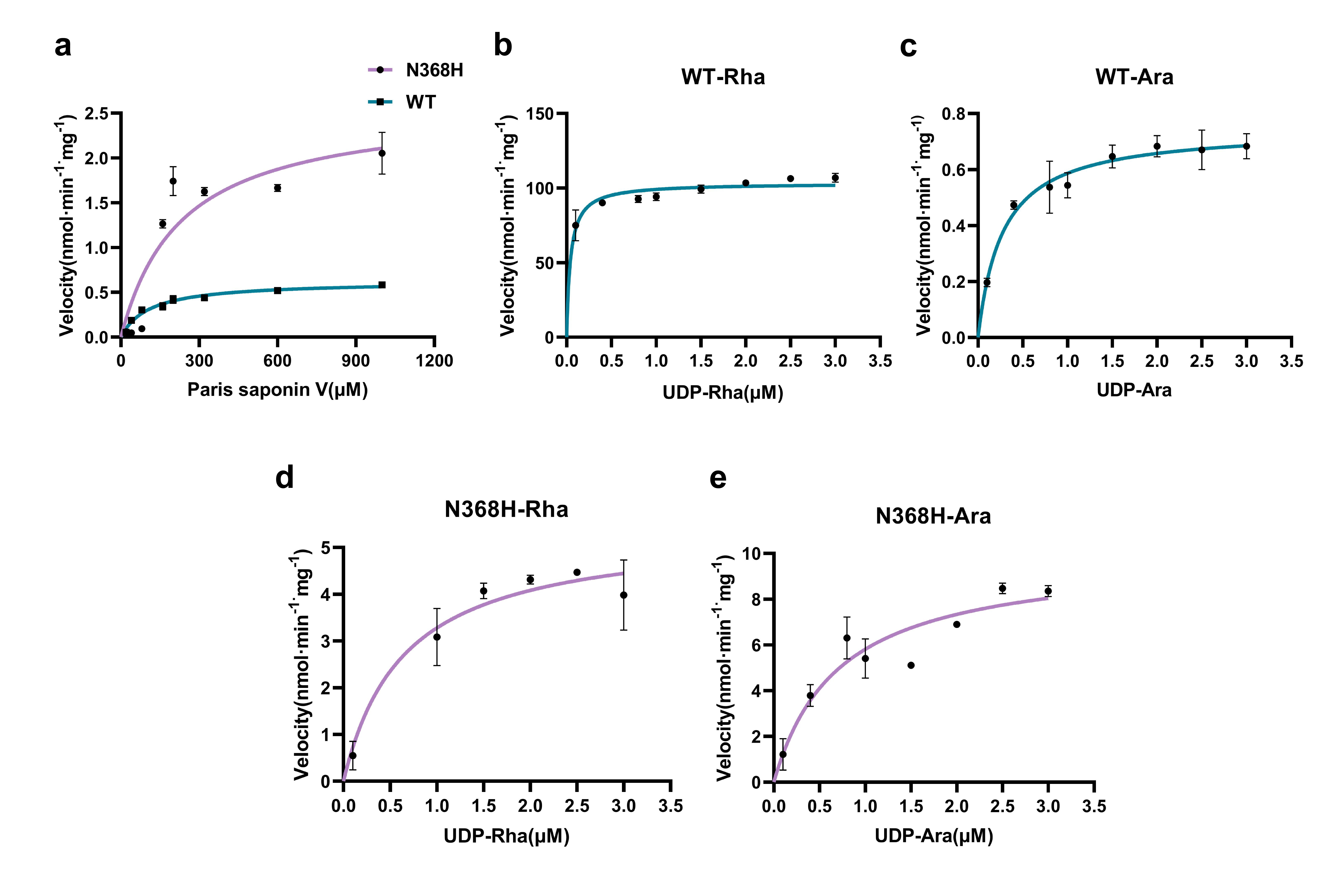

Figure S 1 The kinetic parameters of UGT93M3 and N368H. (a) The Michaelis constant (*K*_m_) of PSⅤ for the WT enzyme and the N368H. (b) The *K*_m_ of UDP-Rha for the WT enzyme. (c) The *K*_m_ of UDP-Ara for the WT enzyme. (d) The *K*_m_ of UDP-Rha for the N368H. (e) The *K*_m_ of UDP-Ara for the N368H.

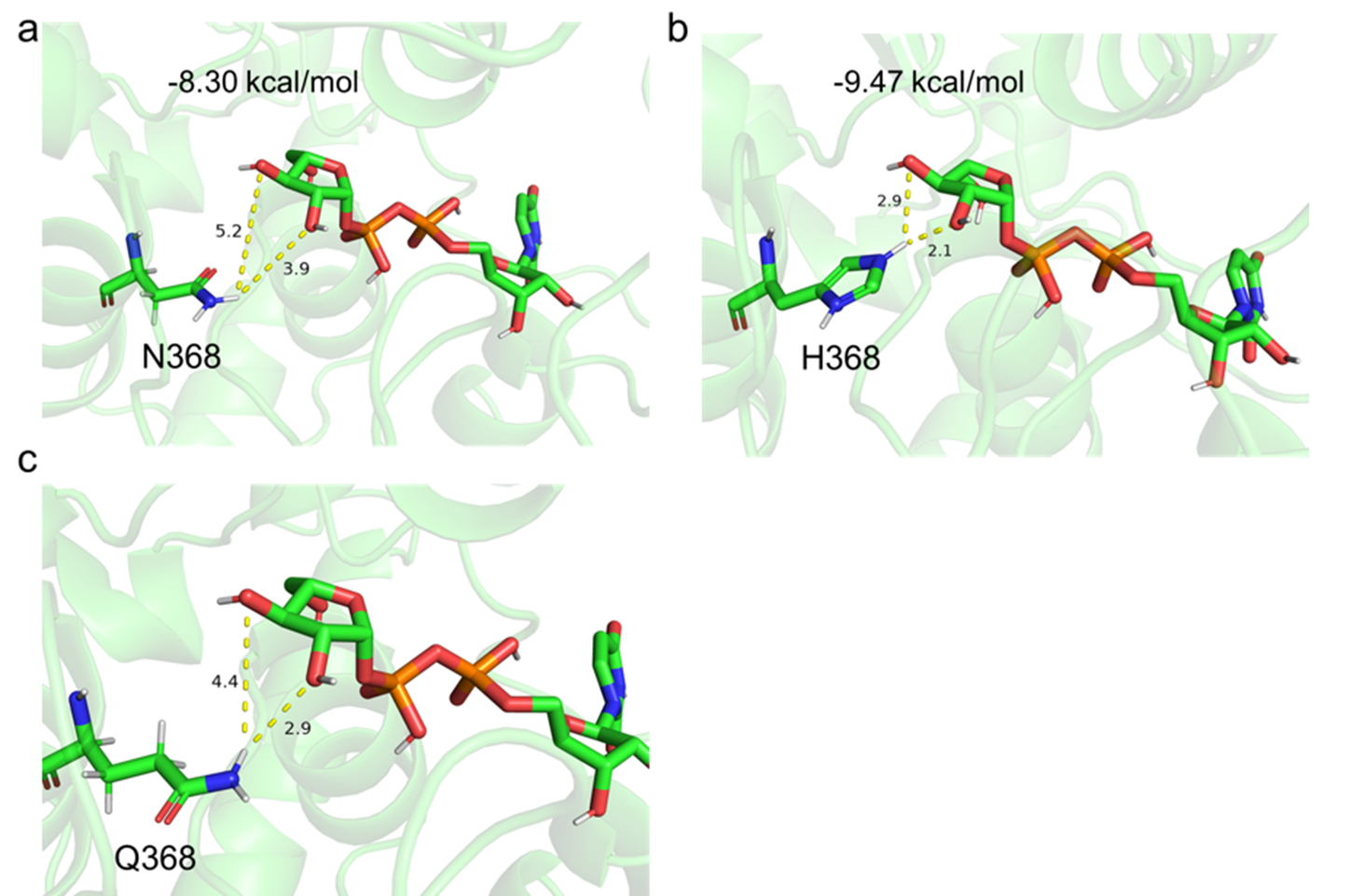

Figure S 2 Molecular docking between N368Q and UDP-Araf revealed that the distances between the terminal hydrogen atom of Q368 and the C3-OH group of UDP-Ara*f* are 2.9 Å and 4.4 Å, respectively.

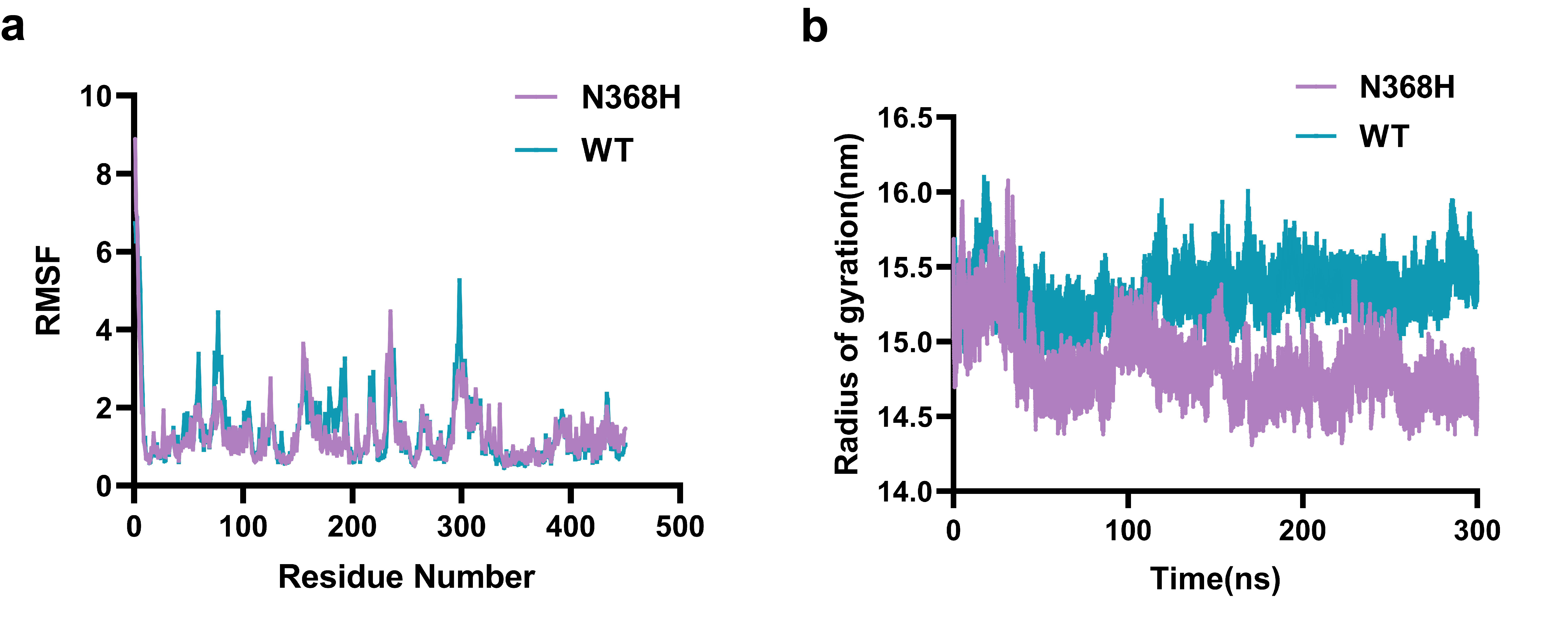

Figure S 3 The molecular dynamics (MD) simulation results of the UGT93M3 enzyme and its N368H mutant. (a) The RMSF analysis reveals that the N368H mutant exhibits enhanced protein structural stability relative to the WT enzyme. (b) The results of radius of gyration (RG) calculation indicate that the N368H mutant has lower protein flexibility.

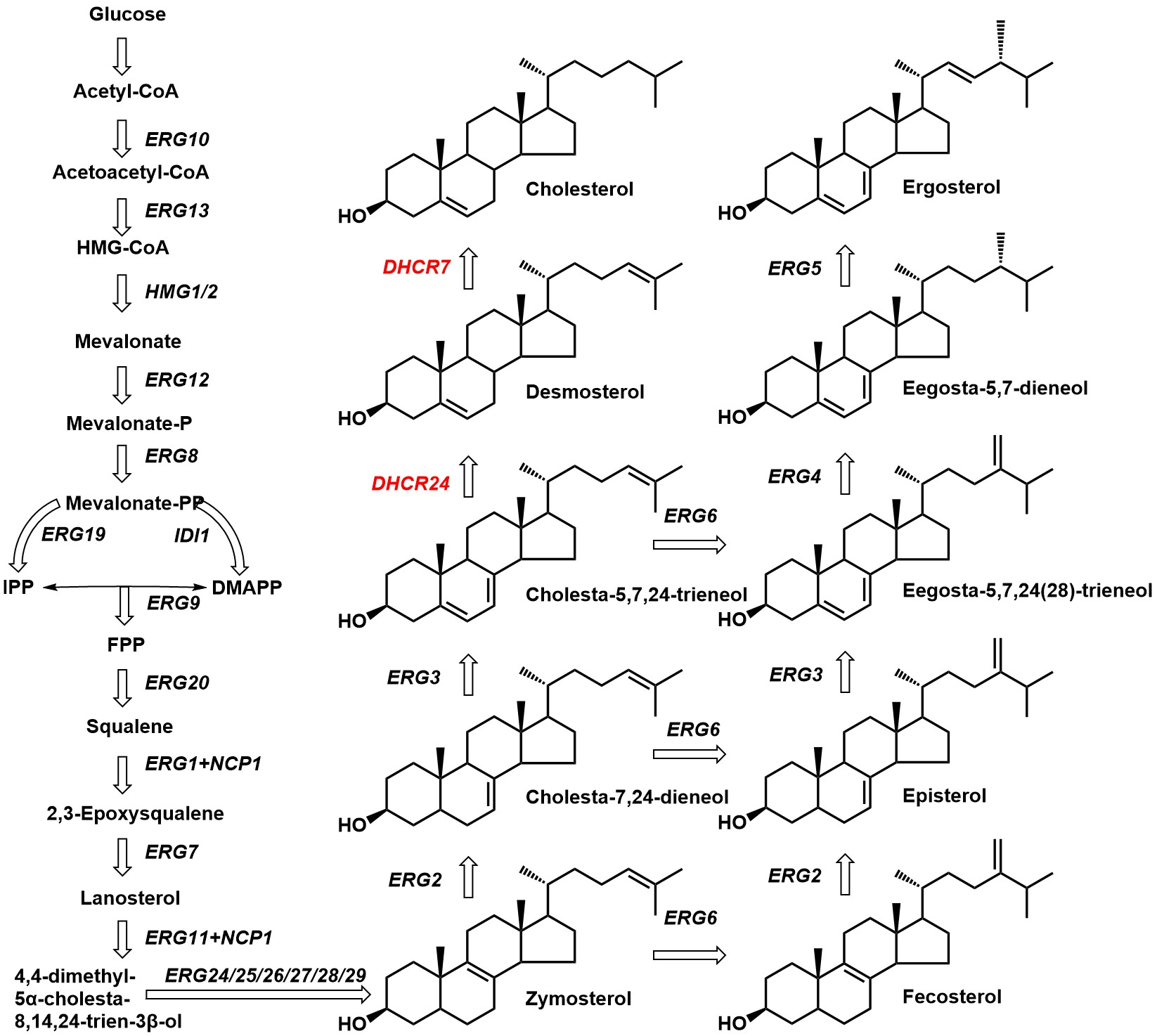

Figure S 4 The synthetic pathways of cholesterol and ergosterol. DHCR7 and DHCR24 are sterol reductases originating from *Solanum tuberosum*, whereas the remaining genes involved are endogenous yeast enzymes. The integration of DHCR7 and DHCR24 enables yeast to synthesize cholesterol. However, the ergosterol biosynthesis pathway harbors a competitive interaction mediated by the ERG6 gene with the cholesterol precursor, which may limit cholesterol accumulation.

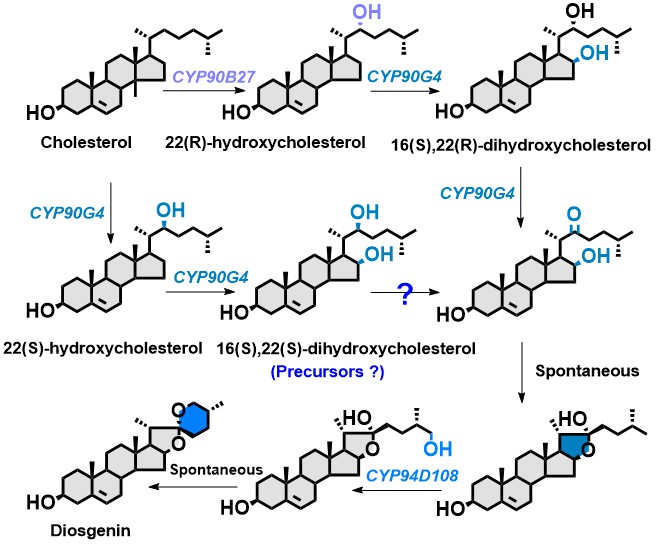

Figure S 5 The multifunctional pathway of CYP90G4. The CYP90B27 gene catalyzes the conversion of cholesterol into 22(*R*)-hydroxycholesterol. This intermediate is subsequently modified by CYP90G4 to produce 16,22-dihydroxycholesterol, which is then converted into diosgenin through the catalytic activity of CYP94D108. In addition, CYP90G4 can also catalyze the hydroxylation of cholesterol to form 22(*S*)-hydroxycholesterol, which is further transformed into 16(*S*),22(*S*)-dihydroxycholesterol. The question mark in the figure indicates that it remains to be determined whether 16(*S*),22(*S*)-dihydroxycholesterol can function as a precursor for diosgenin biosynthesis.

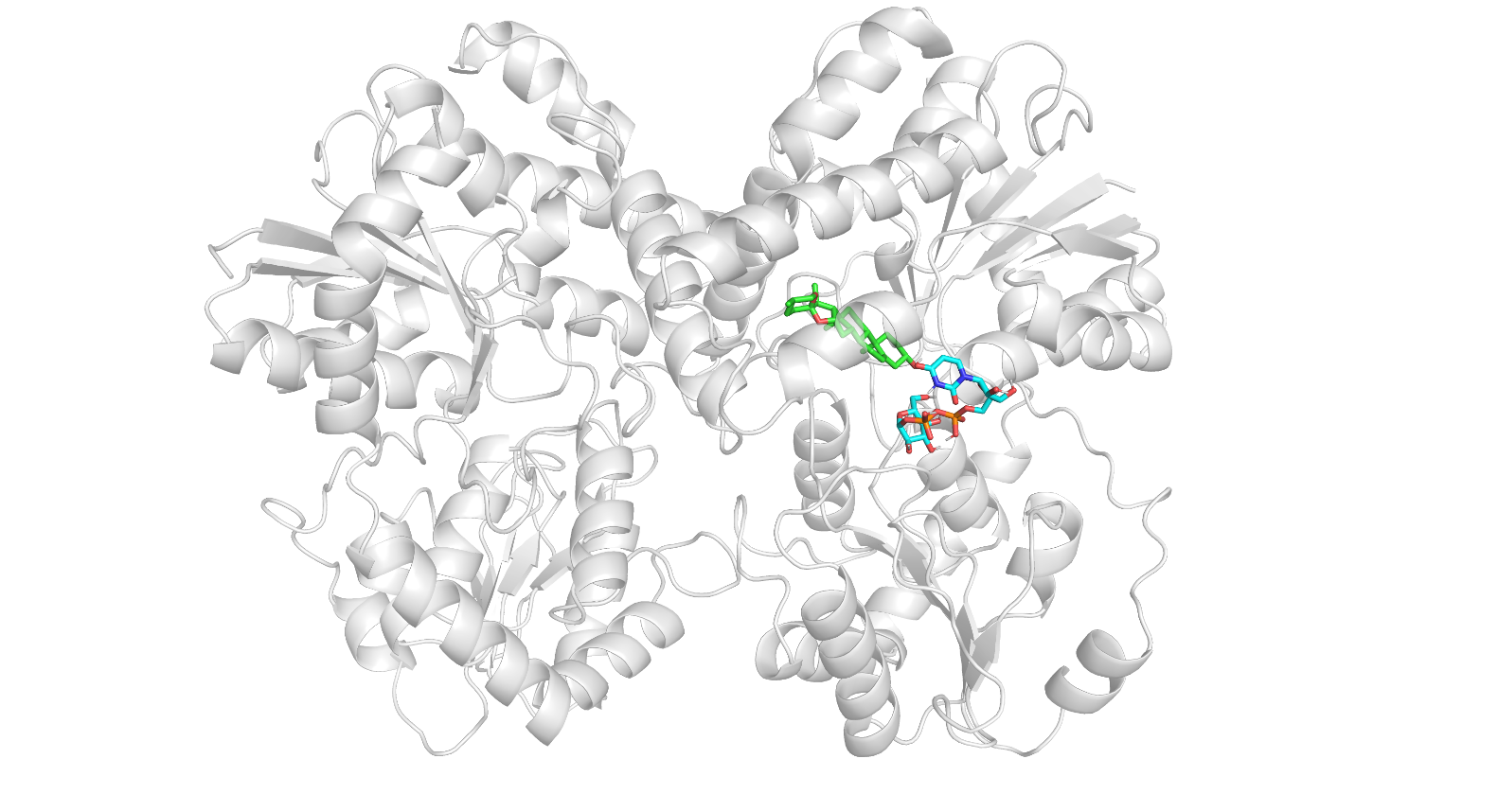

Figure S 6 The AlphaFold3 construction and molecular docking results of *Ppc*UGT4. Molecular docking between the protein receptor and small-molecule ligands (trillin and UDP-Glc) was carried out using AutoDock Vina, and the resulting binding modes were analyzed with Pymol.

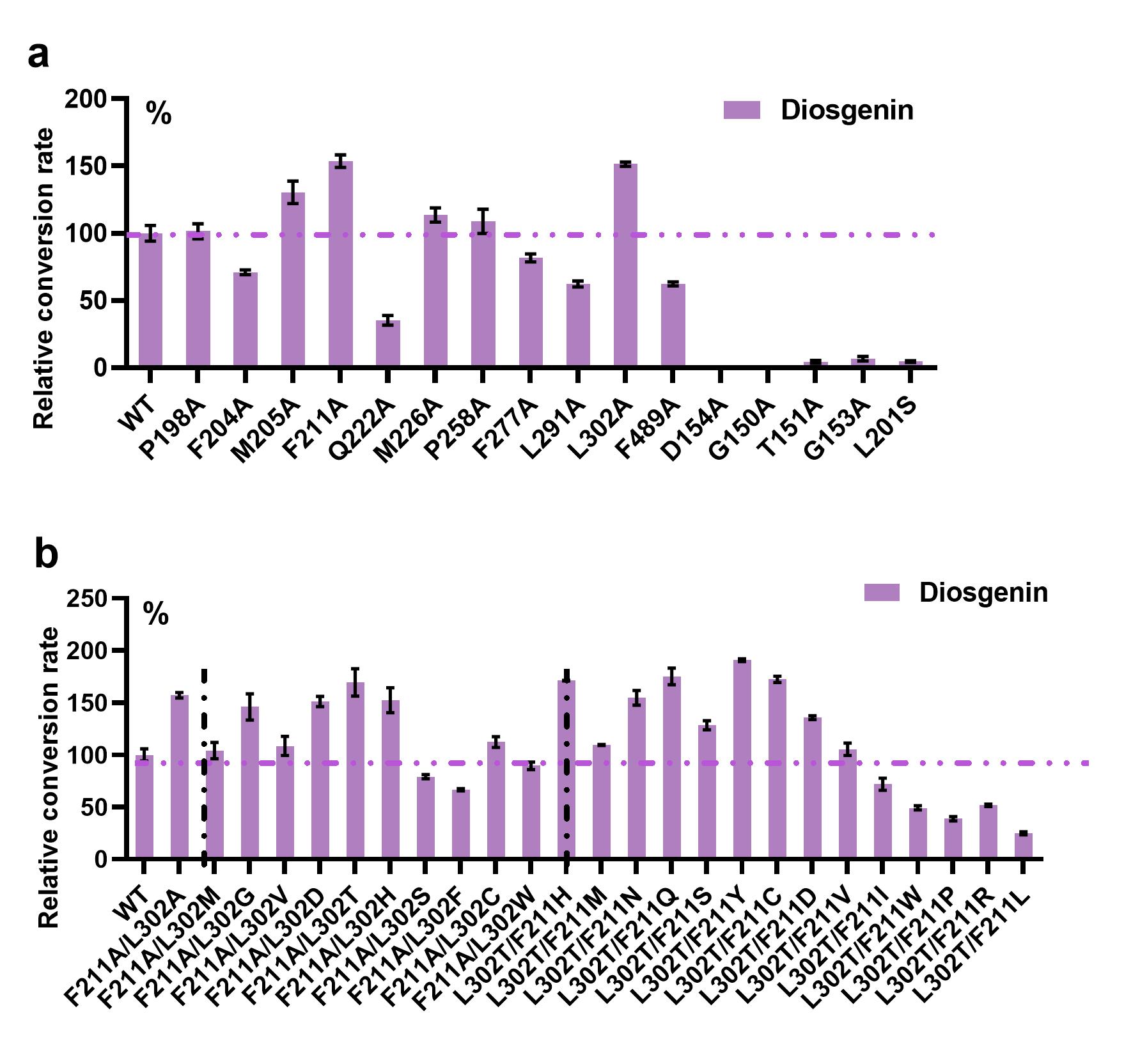

Figure S 7 enzyme engineering modification of *Ppc*UGT4. (a) The relative conversion rate of alanine scanning mutants with diosgenin as the substrate. (b) When iterative mutations were made at the L302 and F211 sites, the relative conversion rate of the mutants using diosgenin as the substrate. All values represent the mean ± SD derived from three biologically independent experiments.

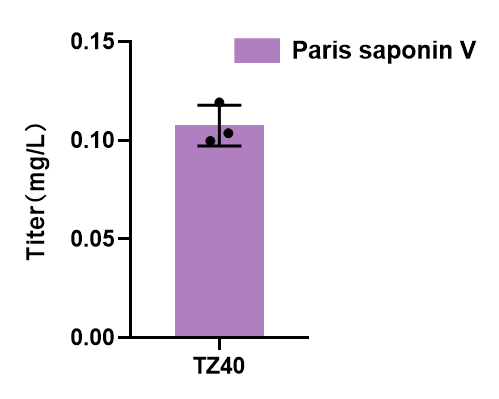

Figure S 8 Fermentation results of the TZ40 strain (PSⅤ production strain). The yeast strains were incubated for 120h at 30 ◦C, 200rpm in YPD medium supplemented with 2 % glucose. All values represent the mean ± SD derived from three biologically independent experiments.

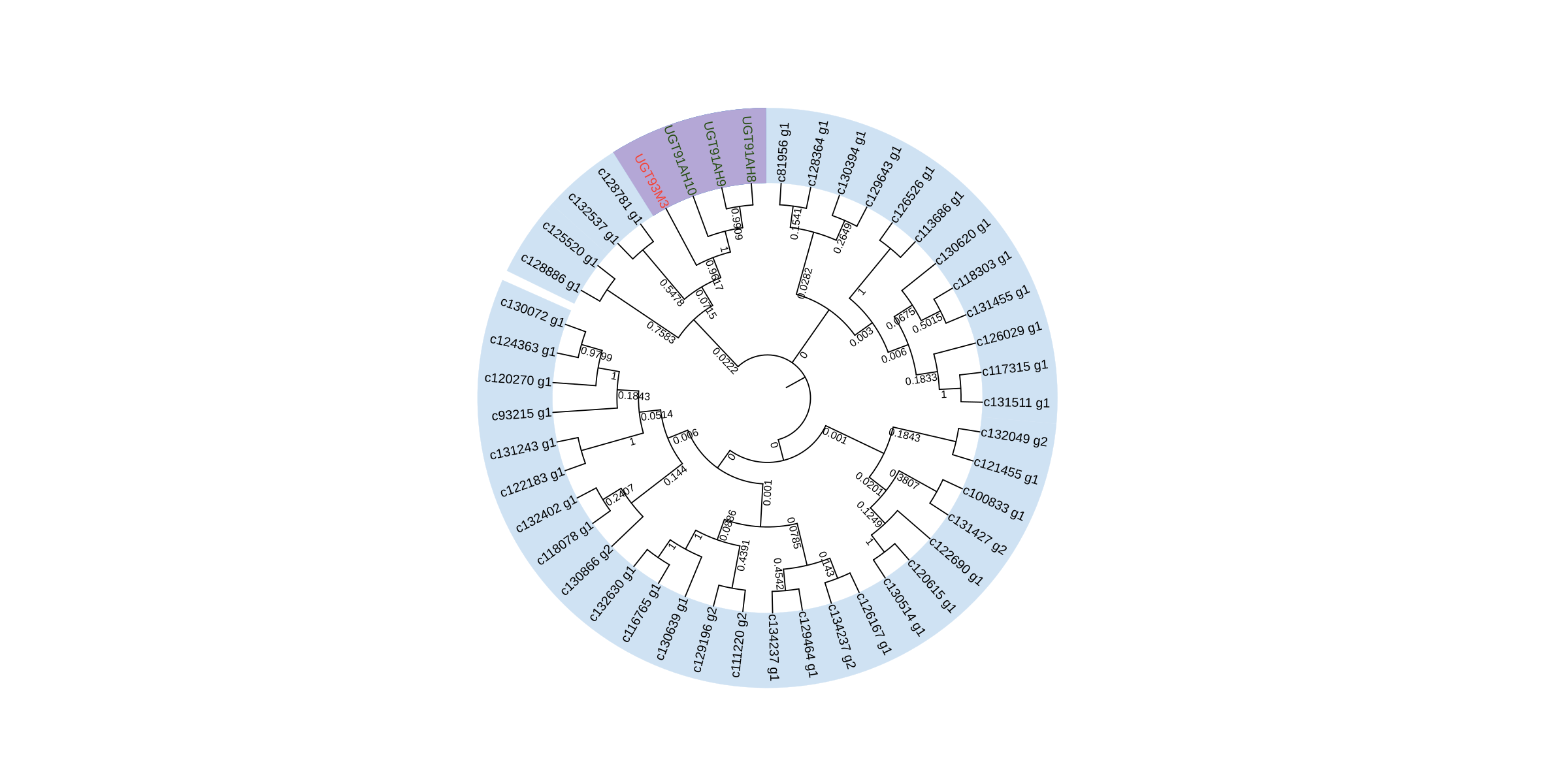

Figure S 9 Phylogenetic analysis of UGT93M3 and transcriptome genes from *P. polyphylla* reveals evolutionary relationships among putative glycosyltransferases. UGT91AH8-10 are phylogenetically closely related to UGT93M3, suggesting a common evolutionary origin and potential functional similarity within the glycosyltransferase family.

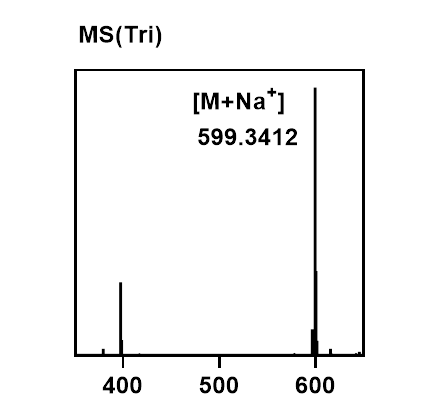

Figure S 10 The characteristic molecular ion peak of trillin

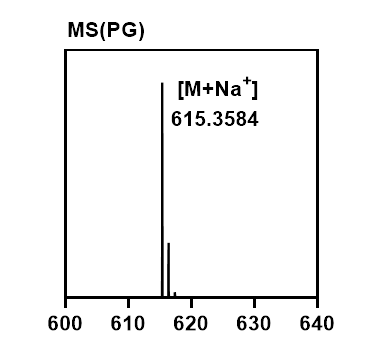

Figure S 11 The characteristic molecular ion peak of Pennogenin-3-O-glucoside

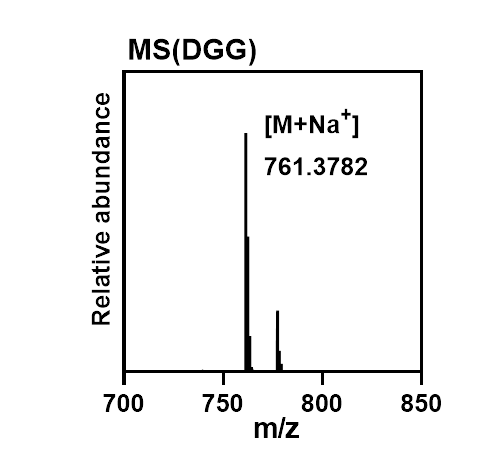

Figure S 12 The characteristic molecular ion peak of Diosgenin-3-O-glucosyl-(1→6)-glucoside (DGG)

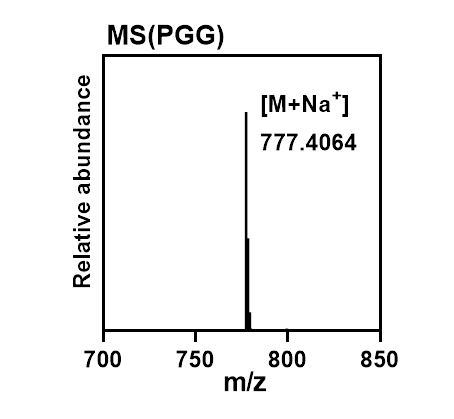

Figure S 13 The characteristic molecular ion peak of Pennogenin-3-O-glucosyl-(1→6)-glucoside (PGG)

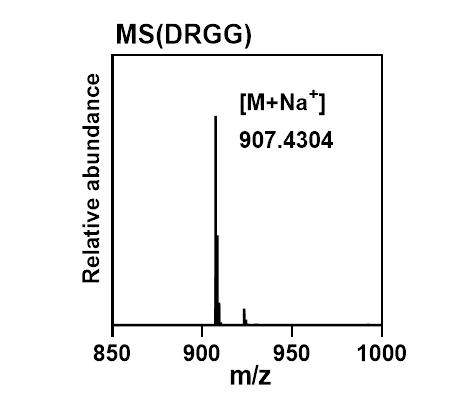

Figure S 14 The characteristic molecular ion peak of Diosgenin-3-O-rhamnosyl(1→2)[glucosyl(1→6)]glucoside (DRGG)

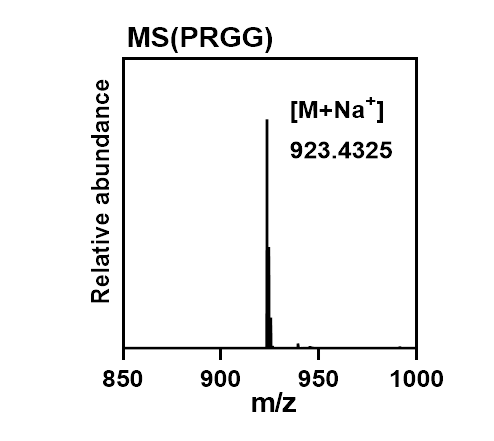

Figure S 15 The characteristic molecular ion peak of Pennogenin-3-O-rhamnosyl(1→2)[glucosyl(1→6)]glucoside (PRGG)

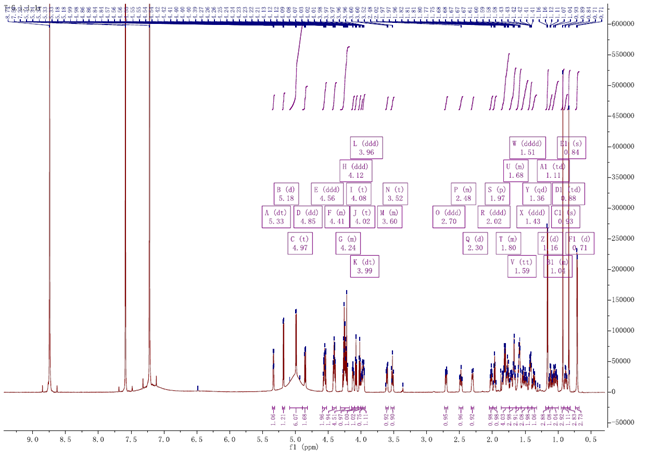

Figure S 16 The ^1^H NMR spectrum of DGG (deuterated pyridine, 800 MHz). δ: 5.33 (dt, J=4.5, 2.1Hz, 1H), 5.18 (d, J=7.8Hz, 1H), 4.99 (d, J=7.8Hz, 2H), 4.85 (dd, J=11.5, 2.0Hz, 1H), 4.59-4.55 (m, 1H), 4.54 (dd, J=9.2, 2.6Hz, 1H), 4.43-4.38 (m, 2H), 4.29-4.25 (m, 2H), 4.25-4.19 (m, 3H), 4.12 (ddd, J=8.7, 5.9, 2.1Hz, 1H), 4.08 (t, J=8.1Hz, 1H), 4.02 (t, J=8.3Hz, 1H), 3.99 (dt, J=6.9, 4.3Hz, 1H), 3.96 (ddd, J=8.3, 5.3, 2.8Hz, 1H), 3.62-3.58 (m, 1H), 3.52 (t, J=10.7Hz, 1H), 2.70 (ddd, J=13.3, 4.8, 2.2Hz, 1H), 2.51-2.45 (m, 1H), 2.30 (d, J=12.7Hz, 1H), 2.02 (ddd, J=12.5, 7.5, 5.6Hz, 1H), 1.96 (p, J=6.9Hz, 1H), 1.86-1.74 (m, 4H), 1.74-1.64 (m, 3H), 1.59 (tt, J=9.3, 5.7Hz, 3H), 1.50 (dddd, J=30.5, 13.9, 10.7, 3.9Hz, 2H), 1.43 (ddd, J=13.6, 11.6, 6.2Hz, 2H), 1.36 (qd, J=13.2, 4.3Hz, 1H), 1.16 (d, J=7.0Hz, 3H), 1.11 (td, J=13.7, 3.7Hz, 1H), 1.09-0.99 (m, 2H), 0.93 (s, 3H), 0.88 (td, J=11.5, 4.7Hz, 1H), 0.84 (s, 3H), 0.71 (d, J=5.6Hz, 3H).

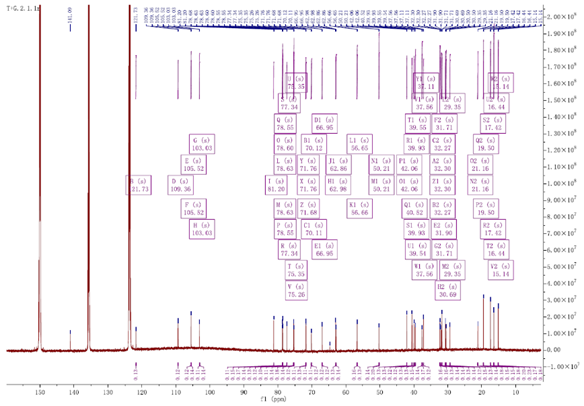

Figure S 17 The ^13^C NMR spectrum of DGG (deuterated pyridine, 200 MHz). δ: C1-C27: 141.58, 122.23, 110.01, 79.04, 70.76, 65.18, 57.15, 50.70, 42.55, 40.42, 40.03, 38.05, 37.60, 32.79, 32.39, 81.69, 63.47, 21.65, 16.93, 19.99, 41.41, 15.63, 31.18, 29.84, 30.99, 67.44, 17.91; 3-Glu: 103.52, 75.75, 79.12, 72.17, 77.83, 70.6; 6’-O-Glu: 106.01, 75.84, 79.09, 72.25, 79.16, 63.35.

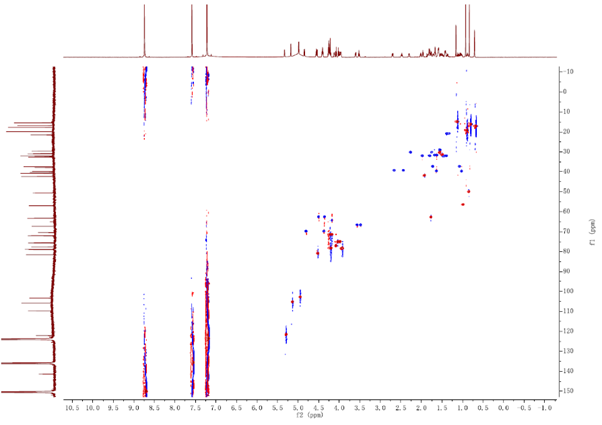

Figure S 18 The HSQC NMR spectrum of DGG

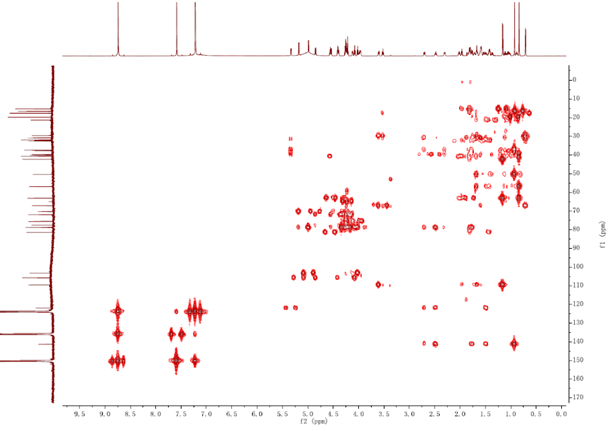

Figure S 19 The HMBC NMR spectrum of DGG

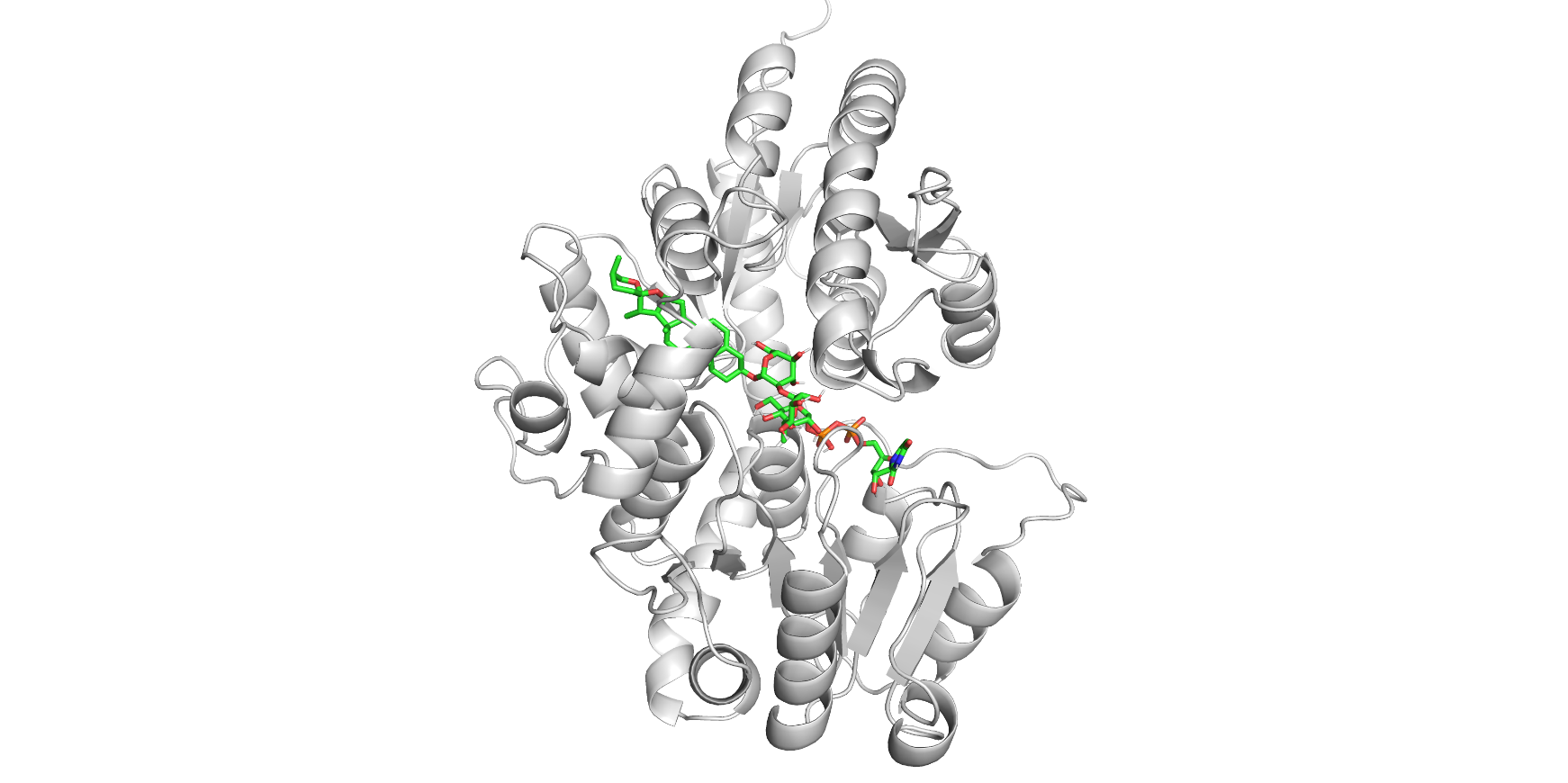

Figure S 20 The AlphaFold 3 modeling and molecular docking results of UGT91AH10. Molecular docking between the protein receptor and small-molecule ligands (PSⅤ and UDP-Glc) was carried out using AutoDock Vina, and the resulting binding modes were analyzed with Pymol.

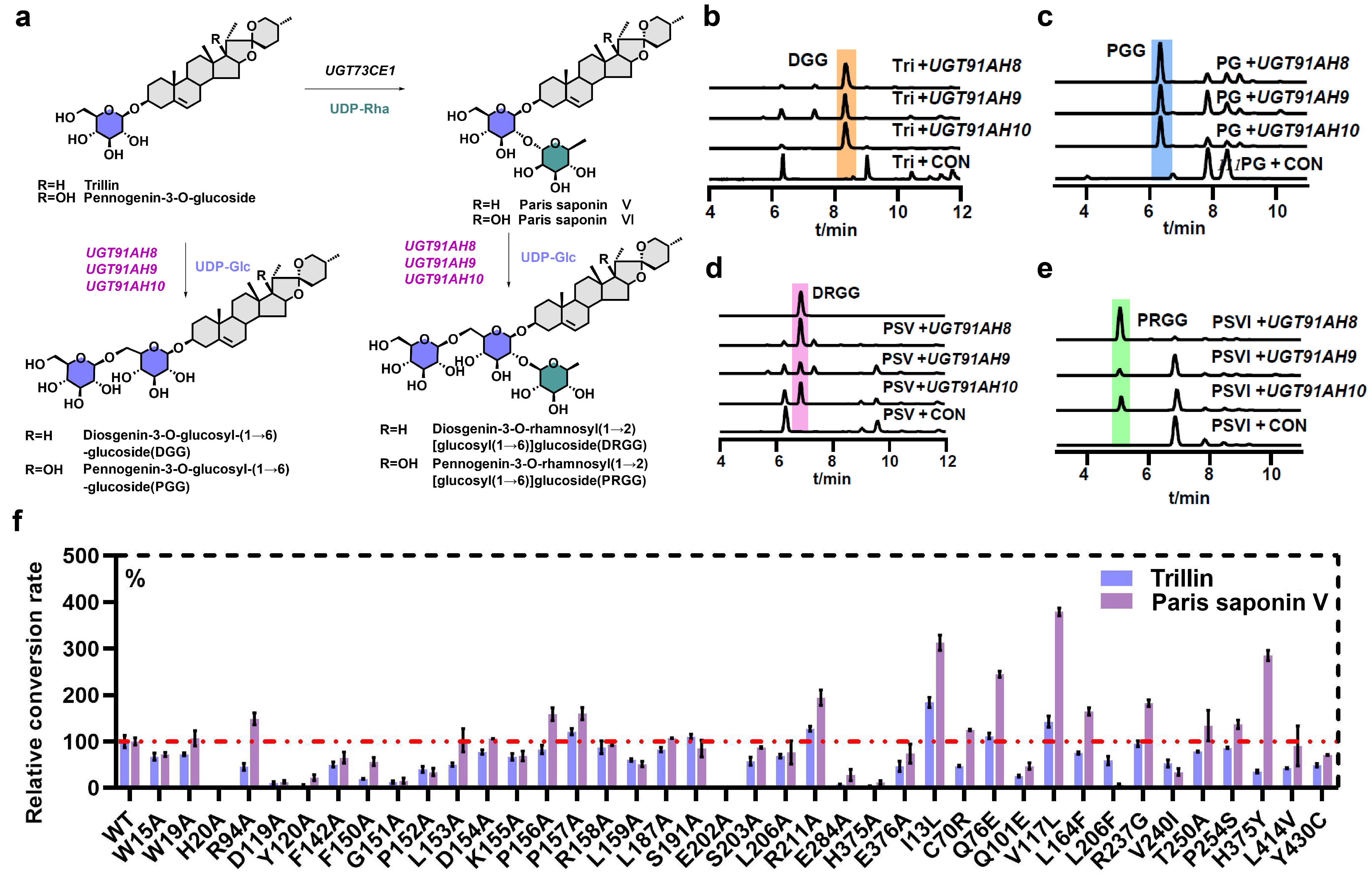

Figure S 21 Functional Characterization of UGT91AH8, UGT91AH9 and UGT91AH10. (a) Catalytic activities of UGT91AH8, UGT91AH9 and UGT91AH10 in saponin biosynthesis, (b) DGG was detected when trillin was employed as the substrate, (c) PGG was detected when PG was employed as the substrate, (d) DRGG was detected when PSV was employed as the substrate. PSV: paris saponin V. (e) PRGG was detected when PSVI was employed as the substrate . PSVI: paris saponin VI. CON: control. (f) In vitro enzyme engineering of UGT91AH10. WT: wild type of UGT91AH10. The nucleotide sequences of UGT91AH8, UGT91AH9 and UGT91AH10 are provided in Supplementary Data 1.

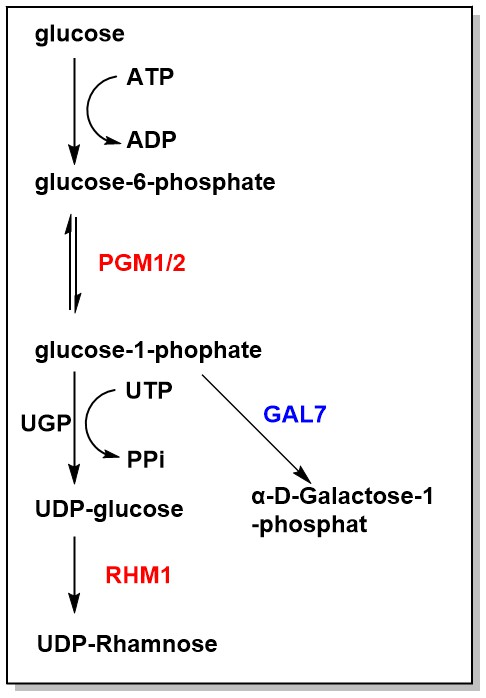

Figure S 22 Modification of the sugar donor pathway. The endogenous yeast gene GAL7 catalyzes the synthesis of α-D-Galactose-1-phosphate, but this process competes with the synthesis of UDP-glucose for precursors.

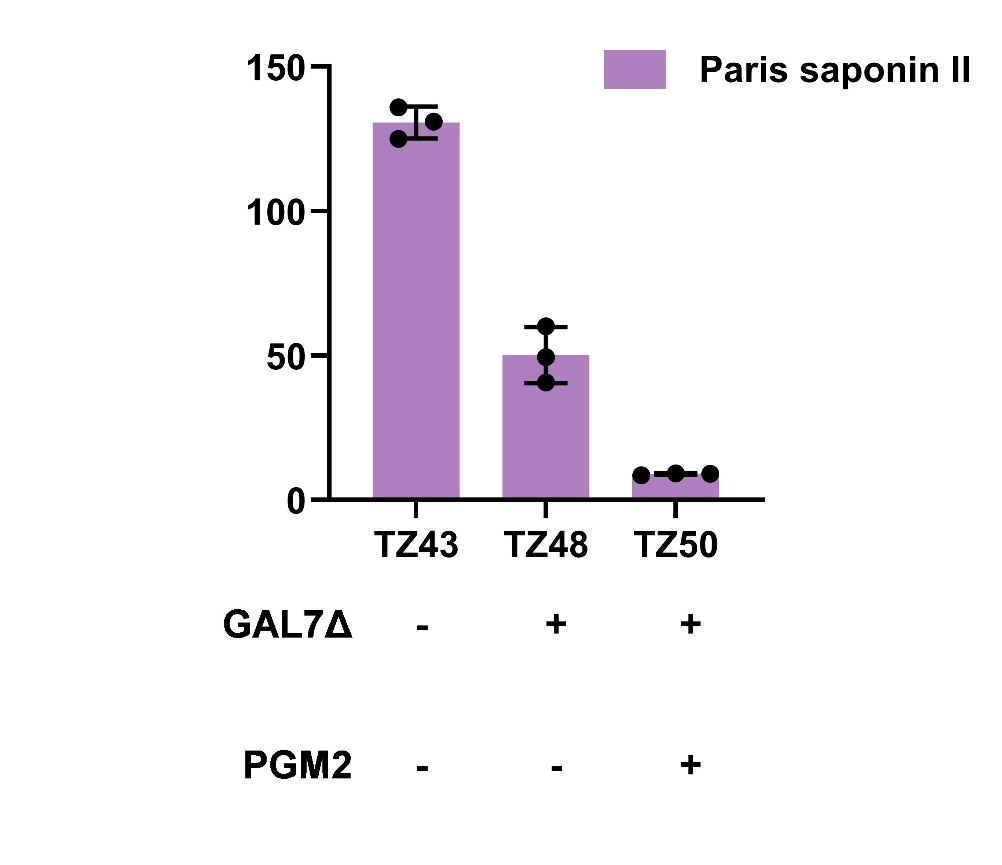

Figure S 23 After the modification of the sugar donor pathway, the yield of the PSⅡ-producing strain decreased. All values represent the mean ± SD derived from three biologically independent experiments.
